## Supplementary Materials for "HAPPY: A deep learning pipeline for mapping cell-to-tissue graphs across placenta histology whole slide images"

**Supplementary Figures 1-7**

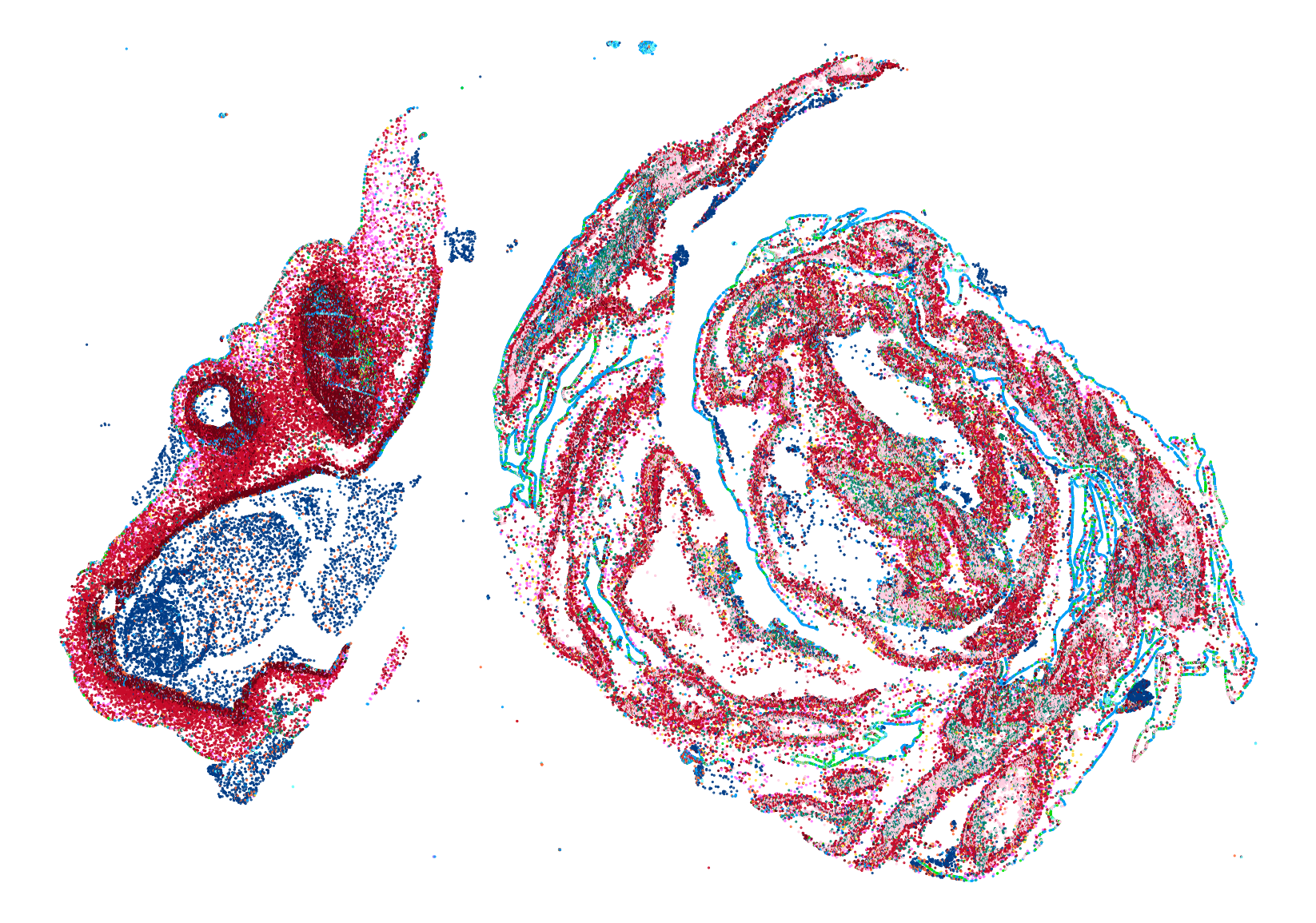

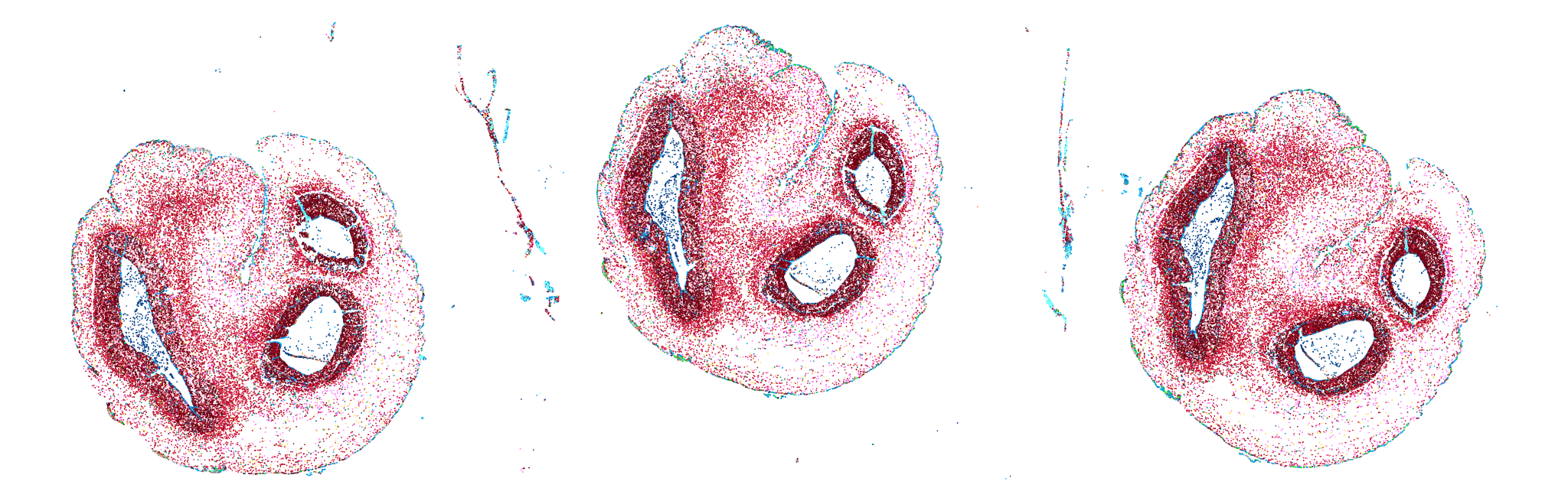

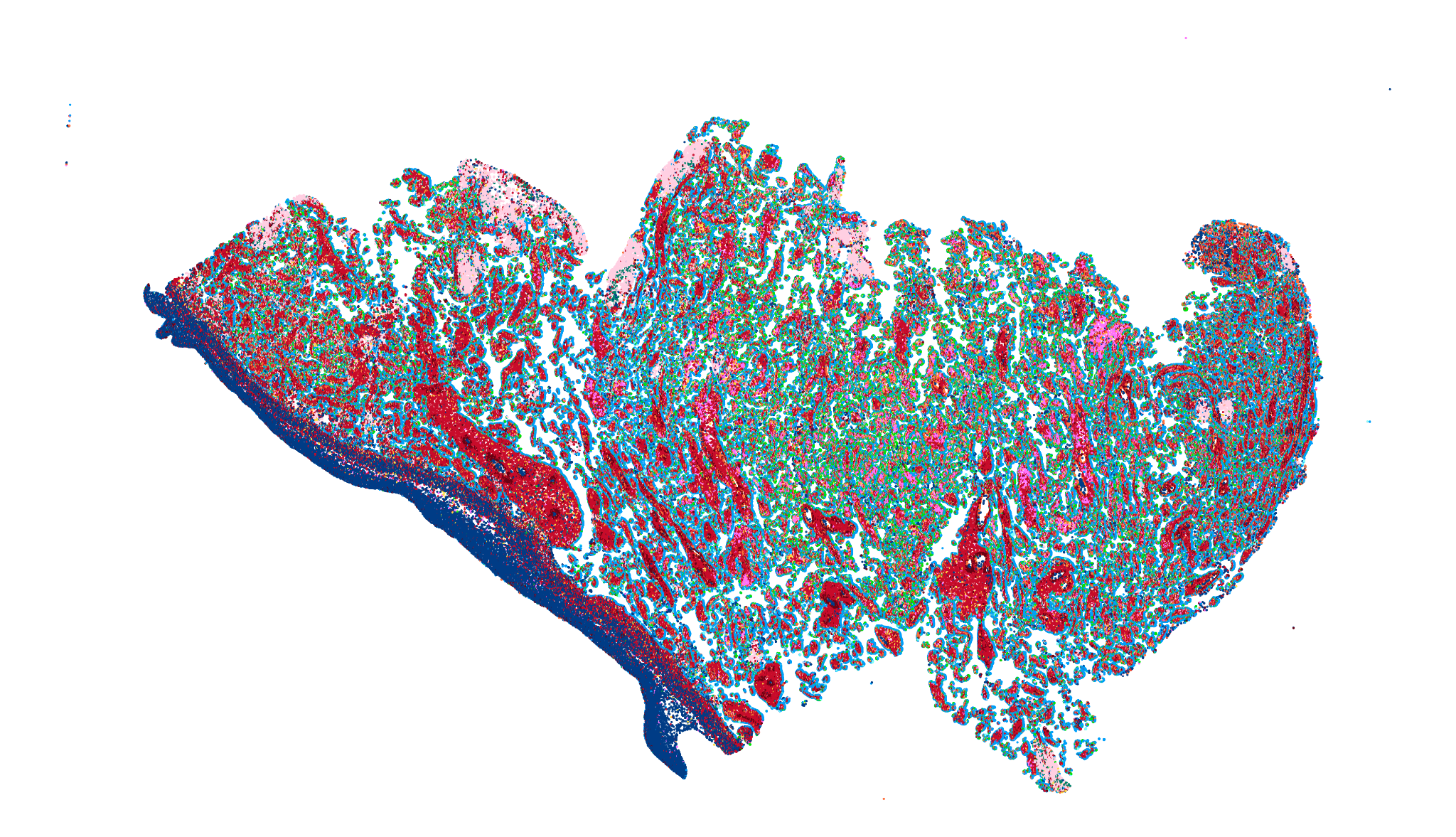

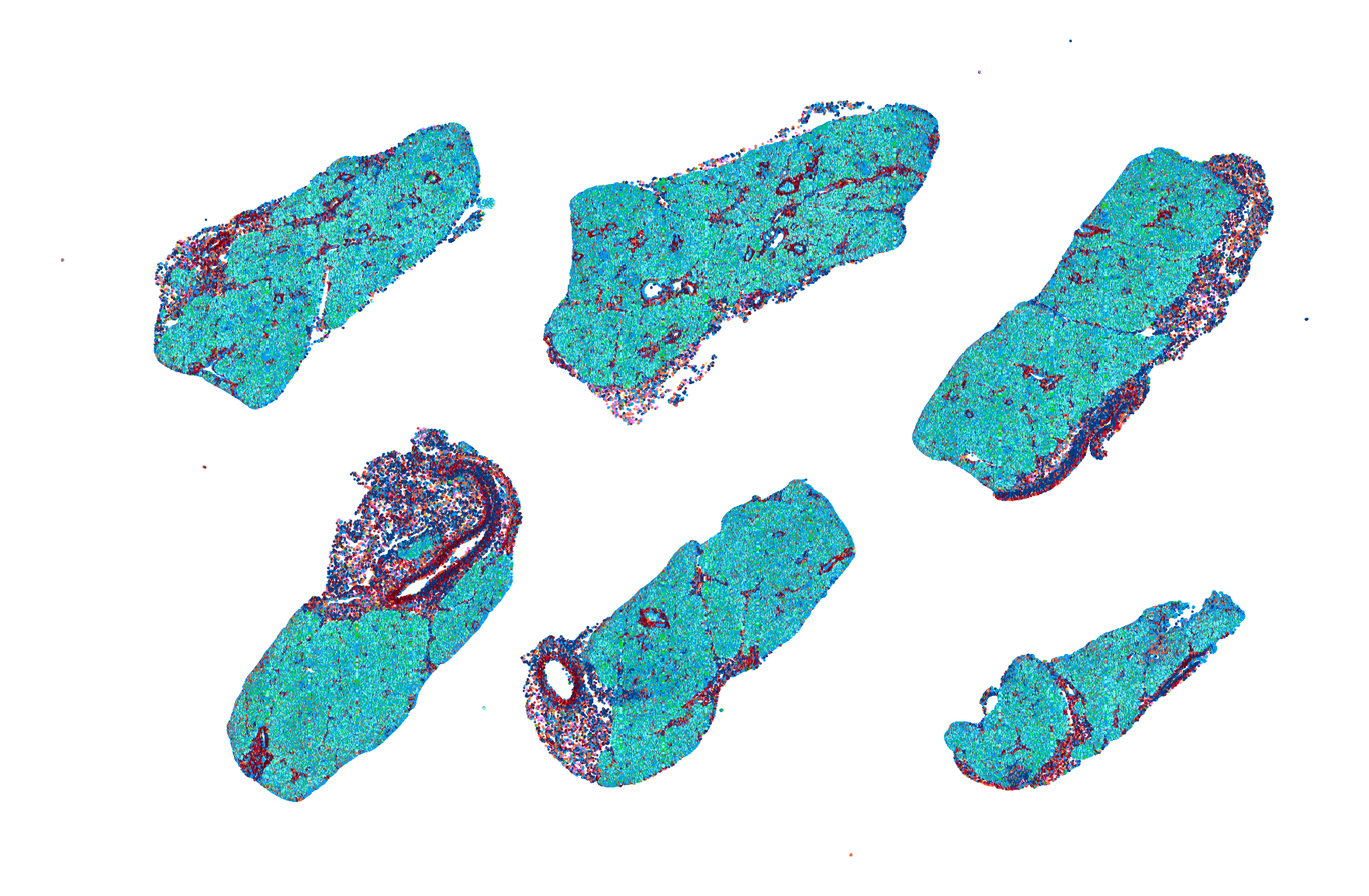

**Supplementary Fig. 1: Visualisation of a preliminary exploration of applying nuclei localisation and cell classification models across other placenta cuts and other organs.** The predictions of nuclei localisation and cell classification models trained for placenta parenchyma histology applied to healthy term placenta membrane rolls (a), healthy term placenta umbilical cords (b), second trimester placental parenchyma with chorioamnionitis (c) from our in-house data and (d) pancreas samples from the GTEx dataset.

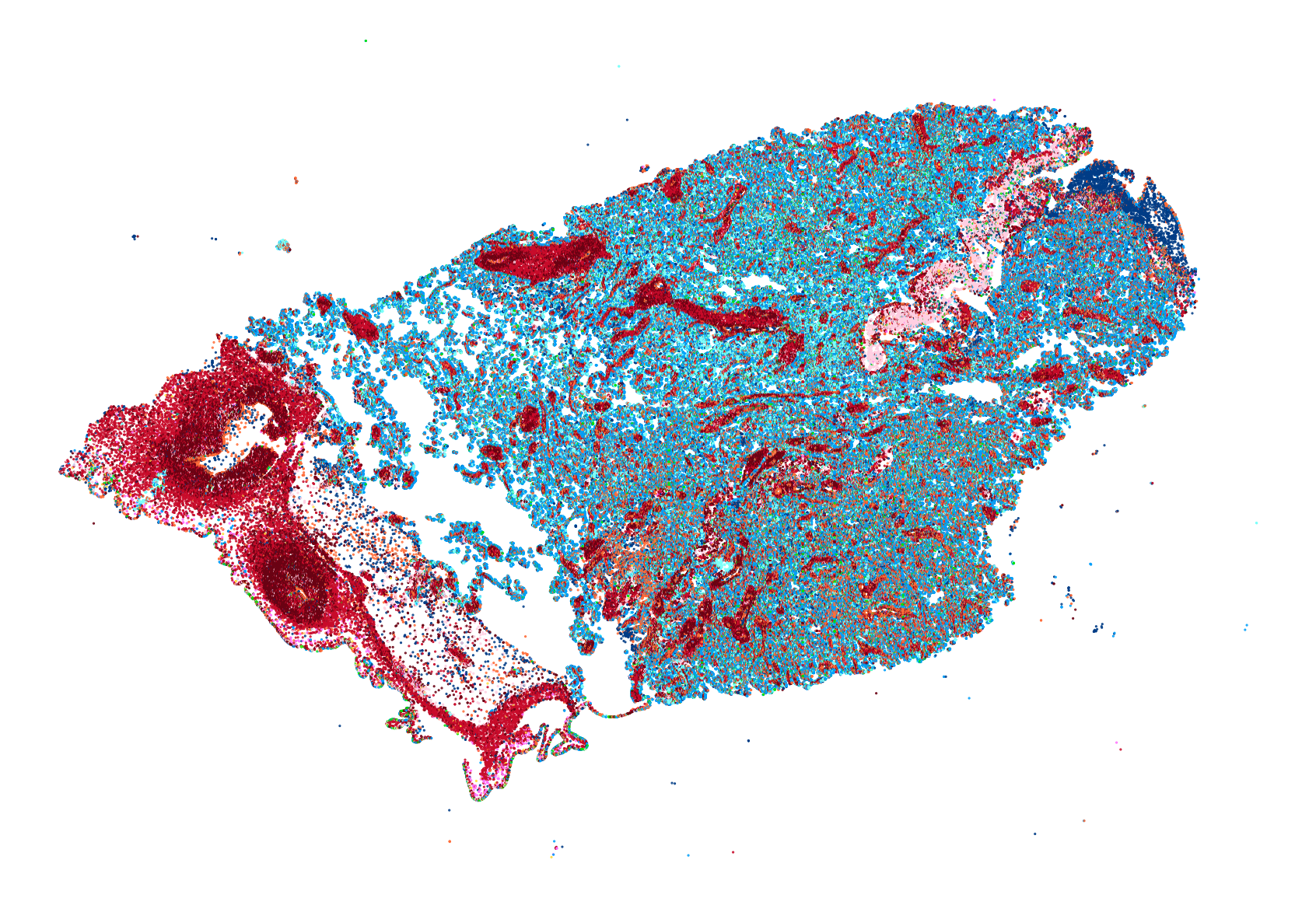

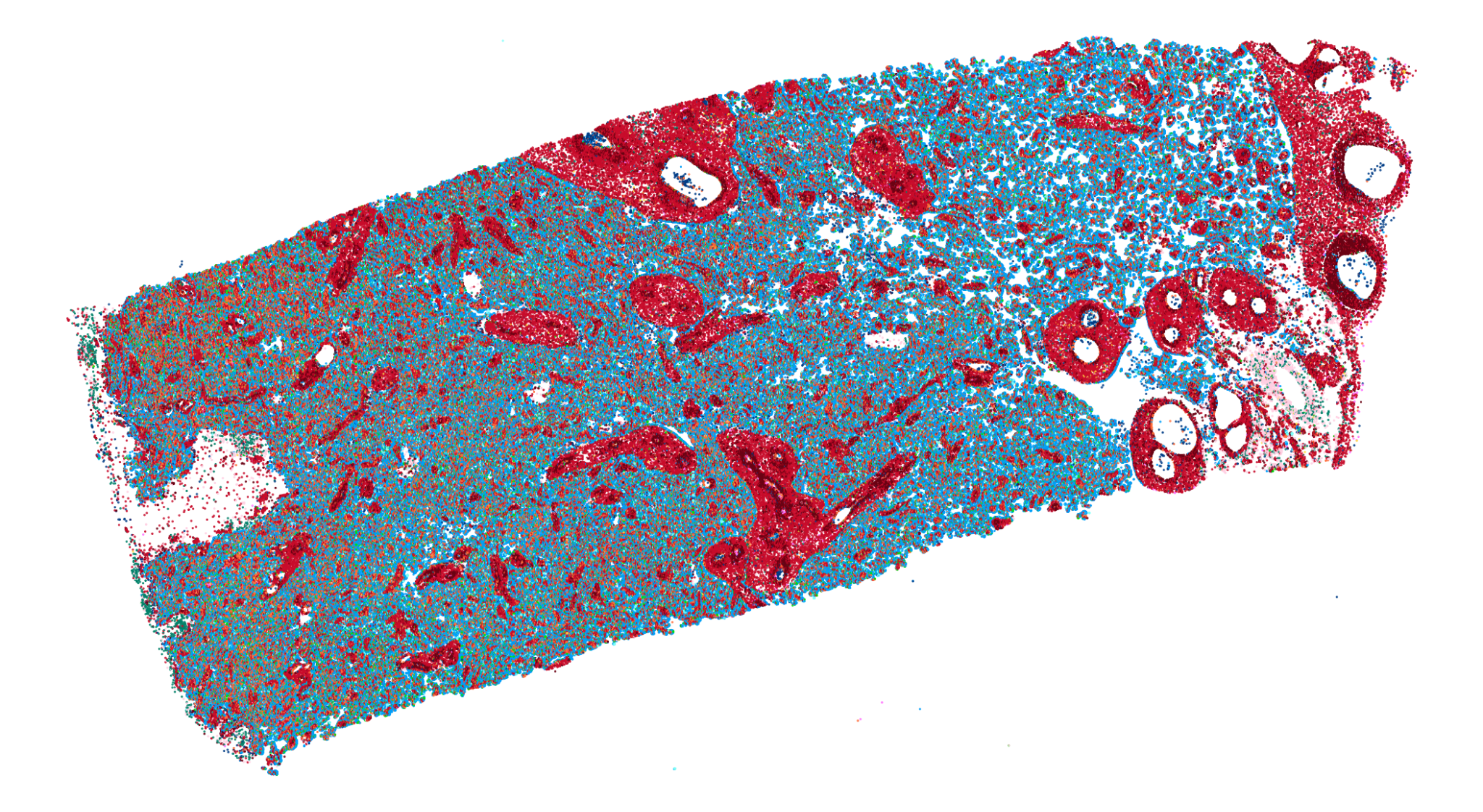

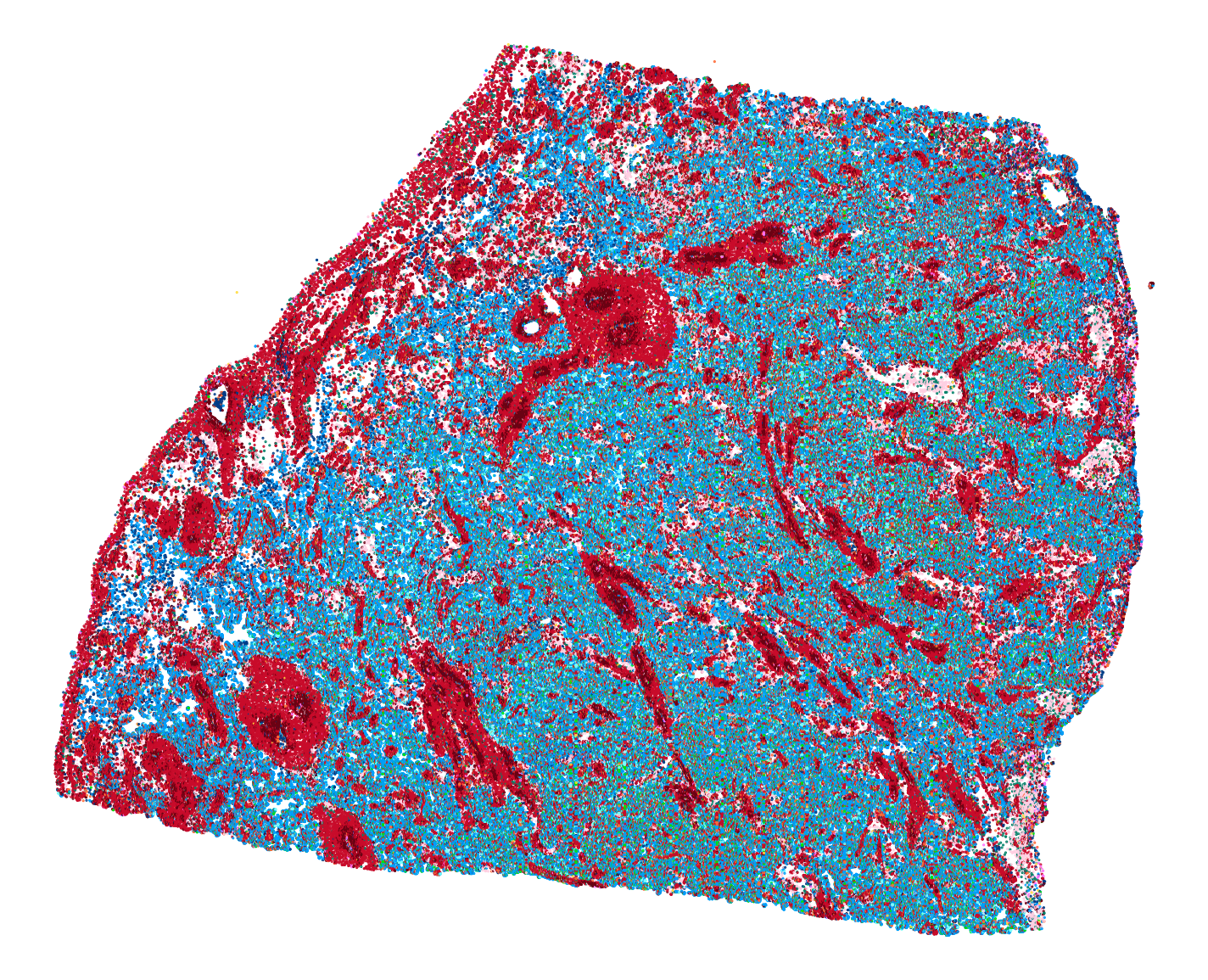

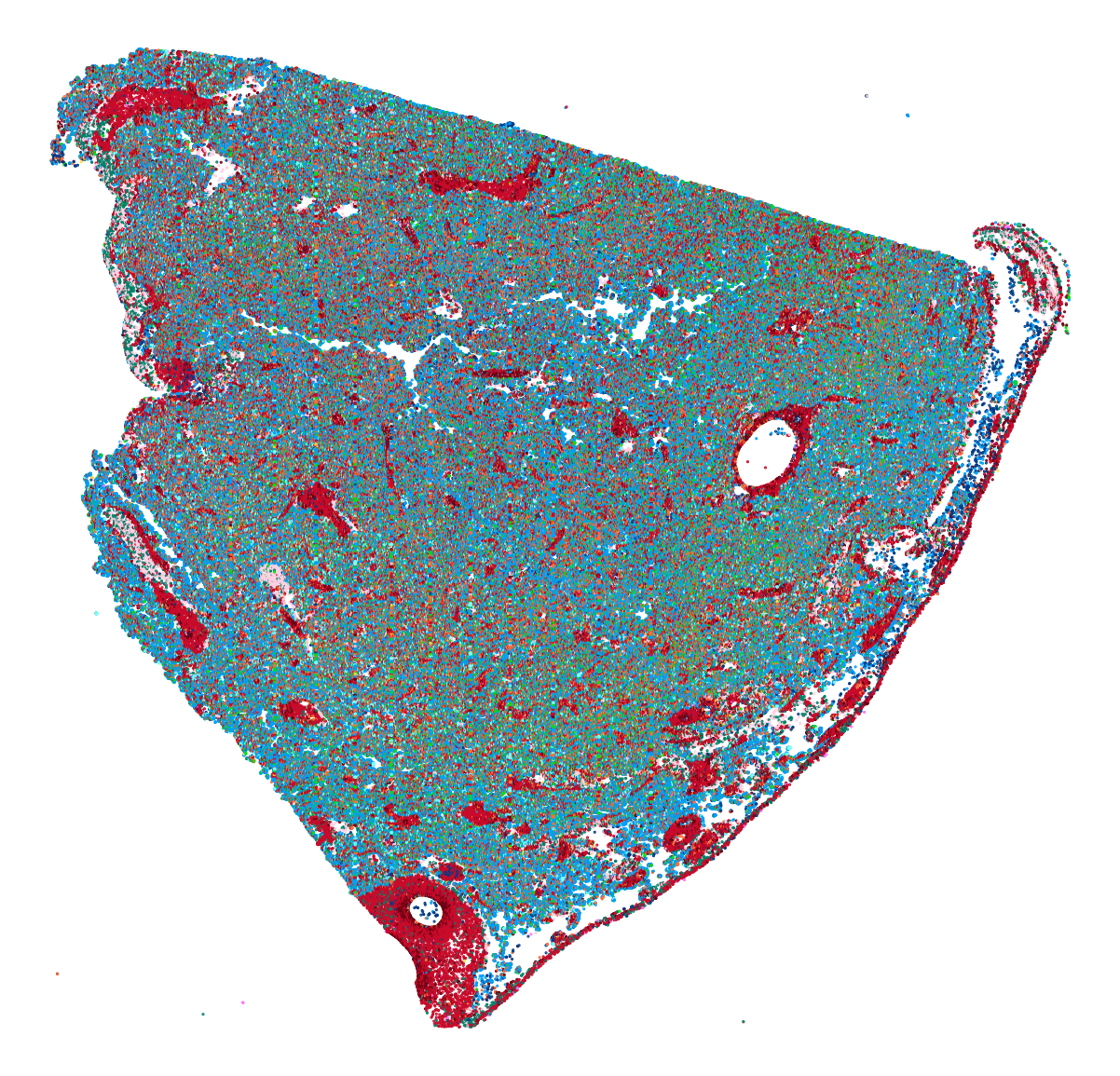

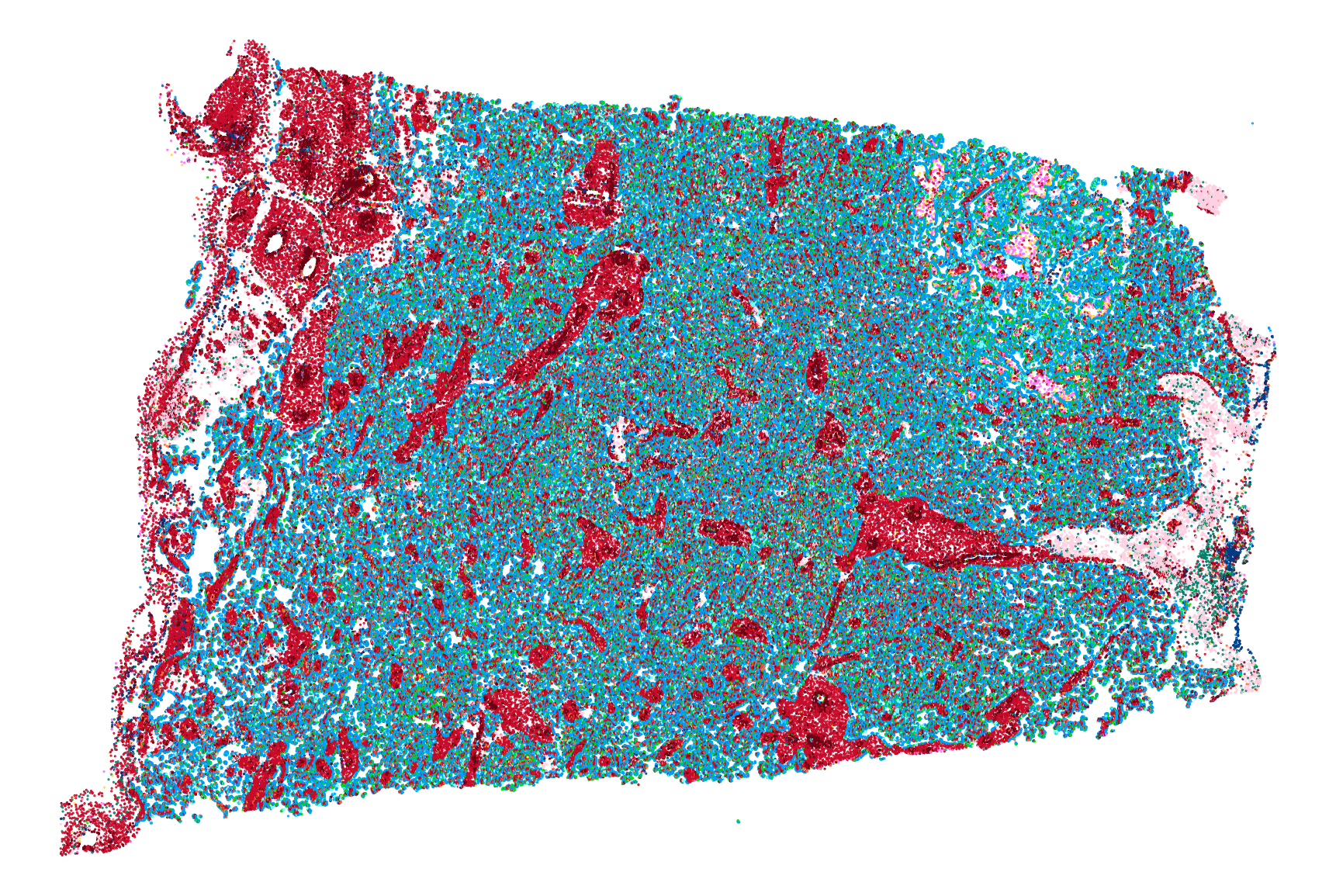

**Supplementary Fig. 2: Visualisation of nuclei and cell predictions across five WSIs of healthy term placenta parenchyma.**

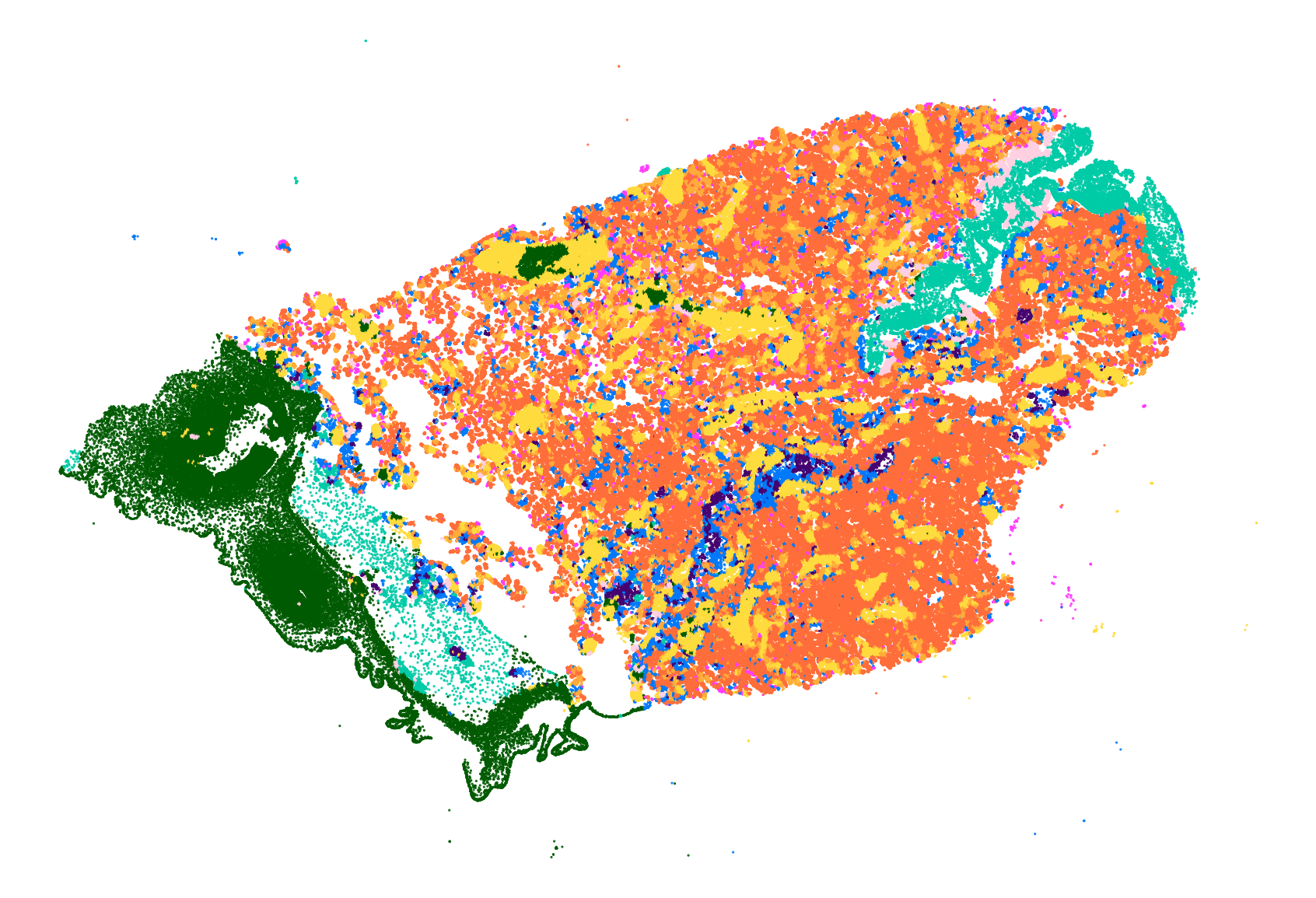

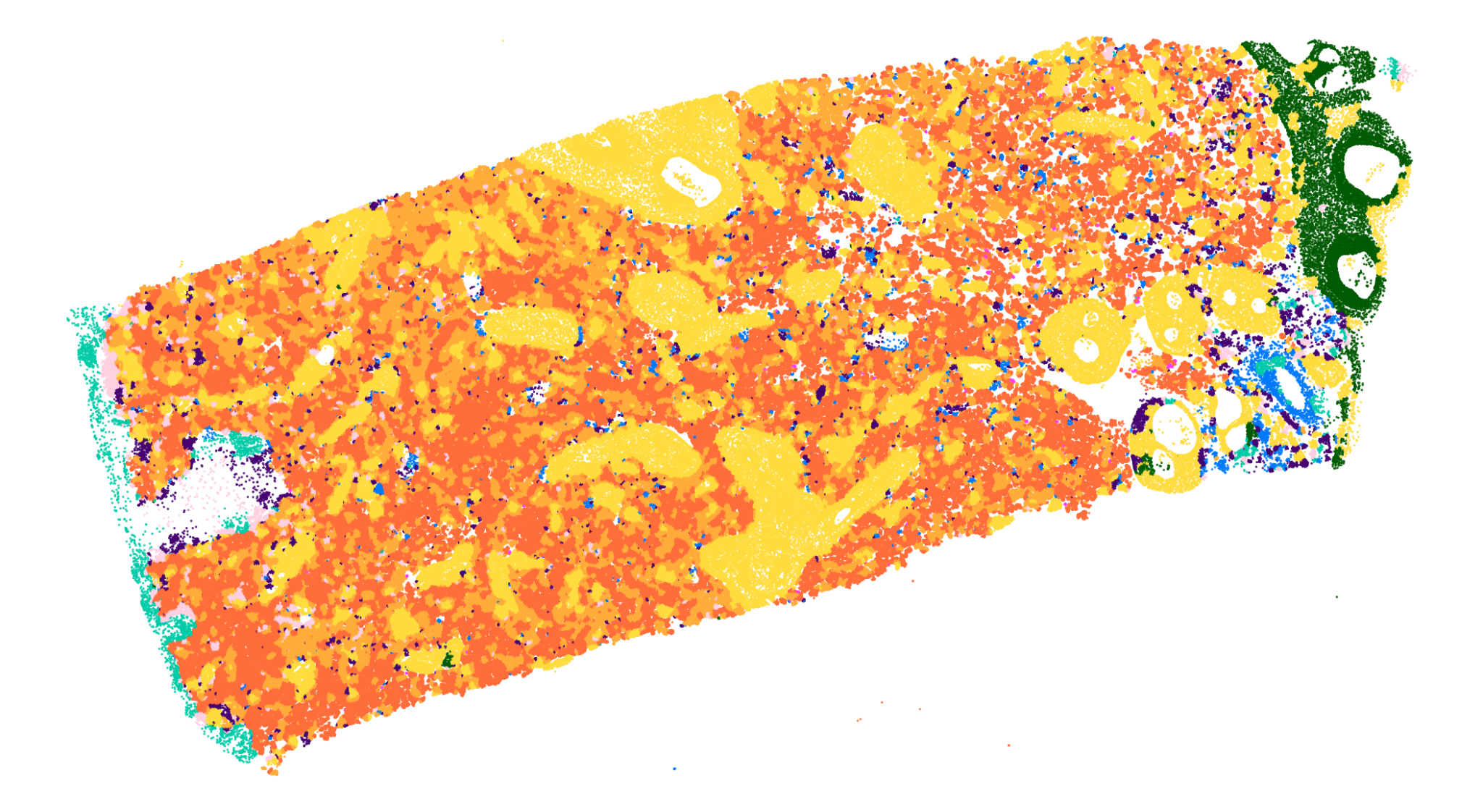

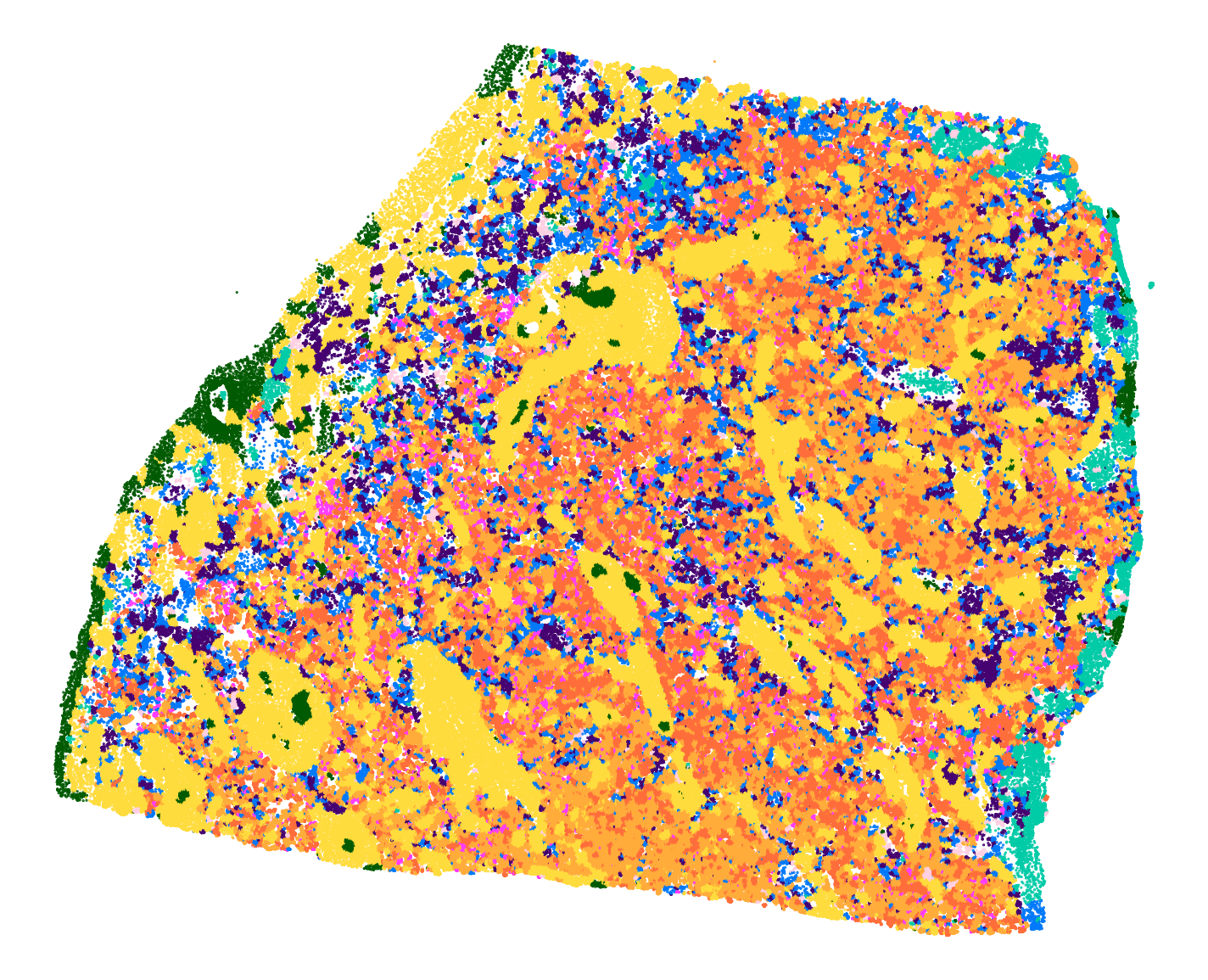

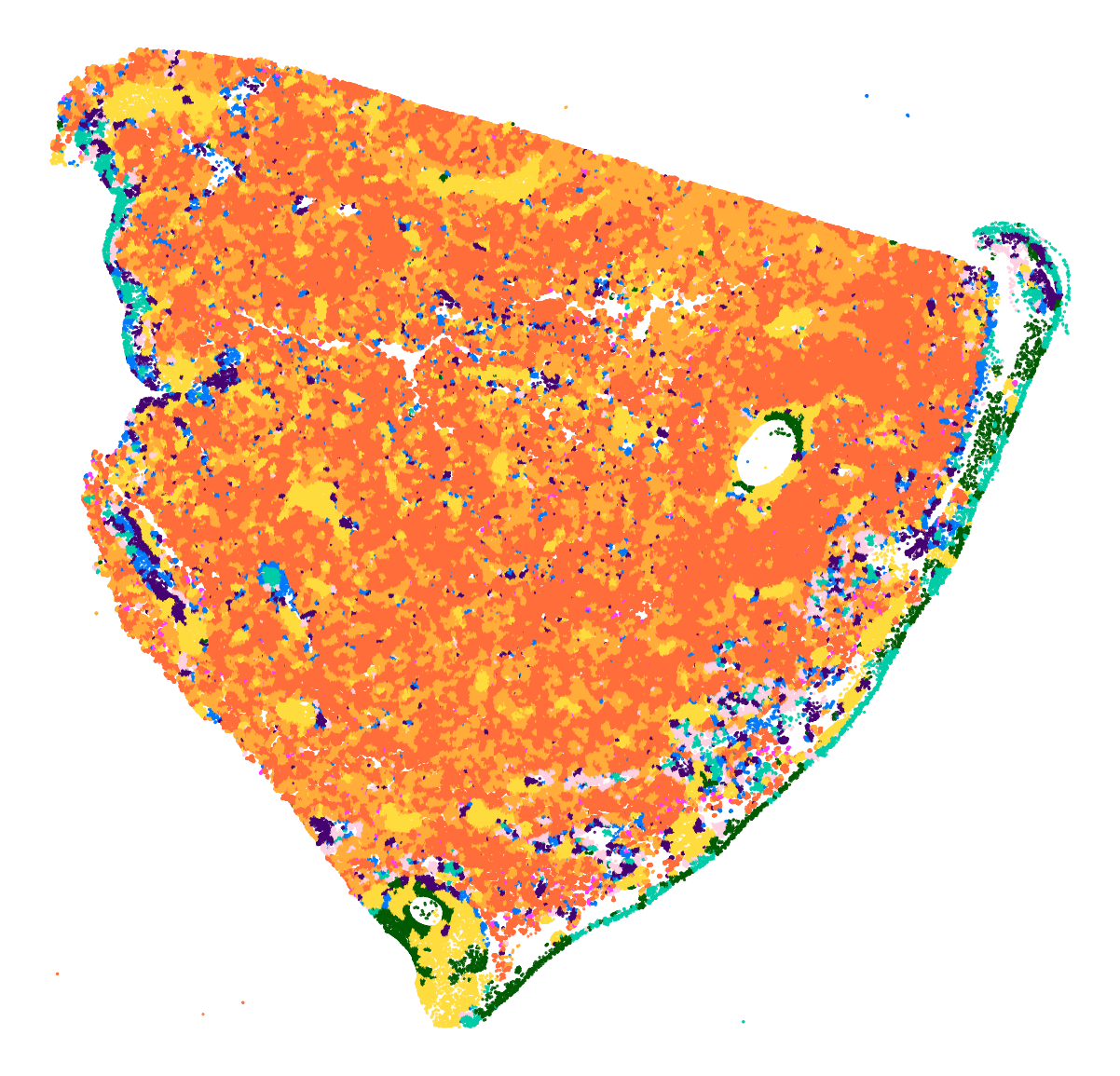

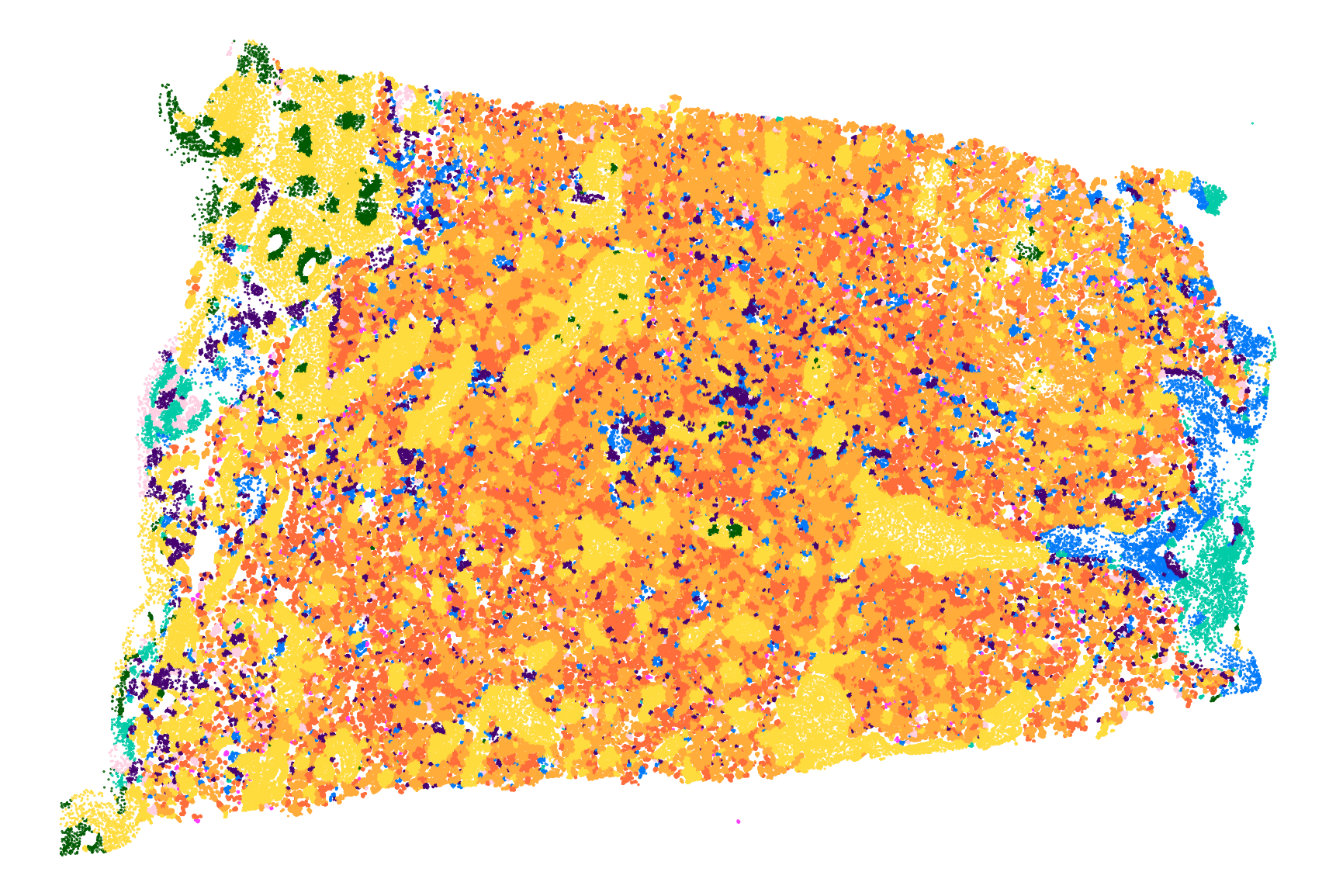

**Supplementary Fig. 3: Visualisation of tissue node predictions across five WSIs of healthy term placenta parenchyma.**

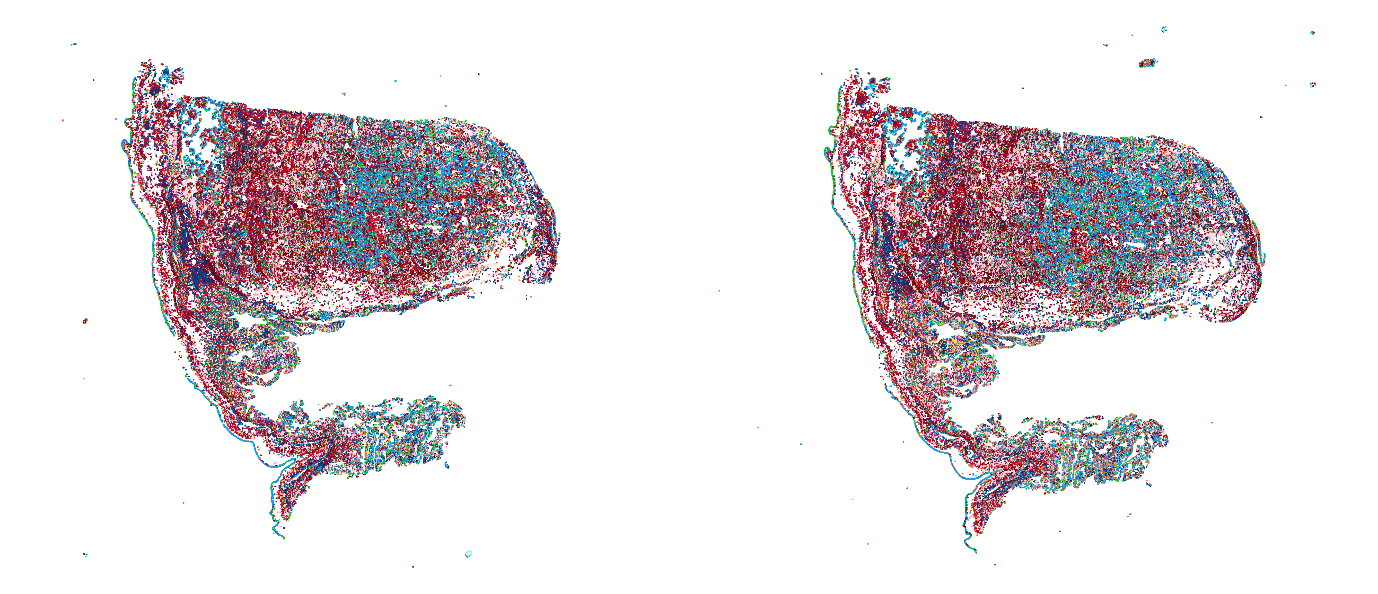

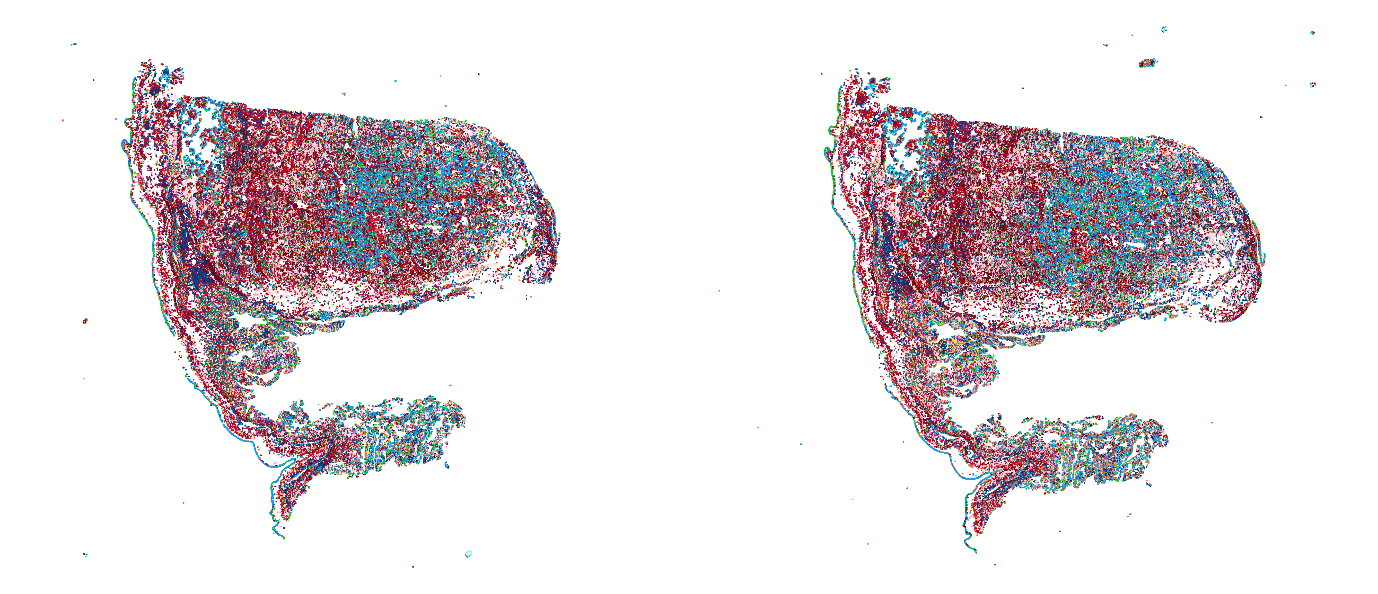

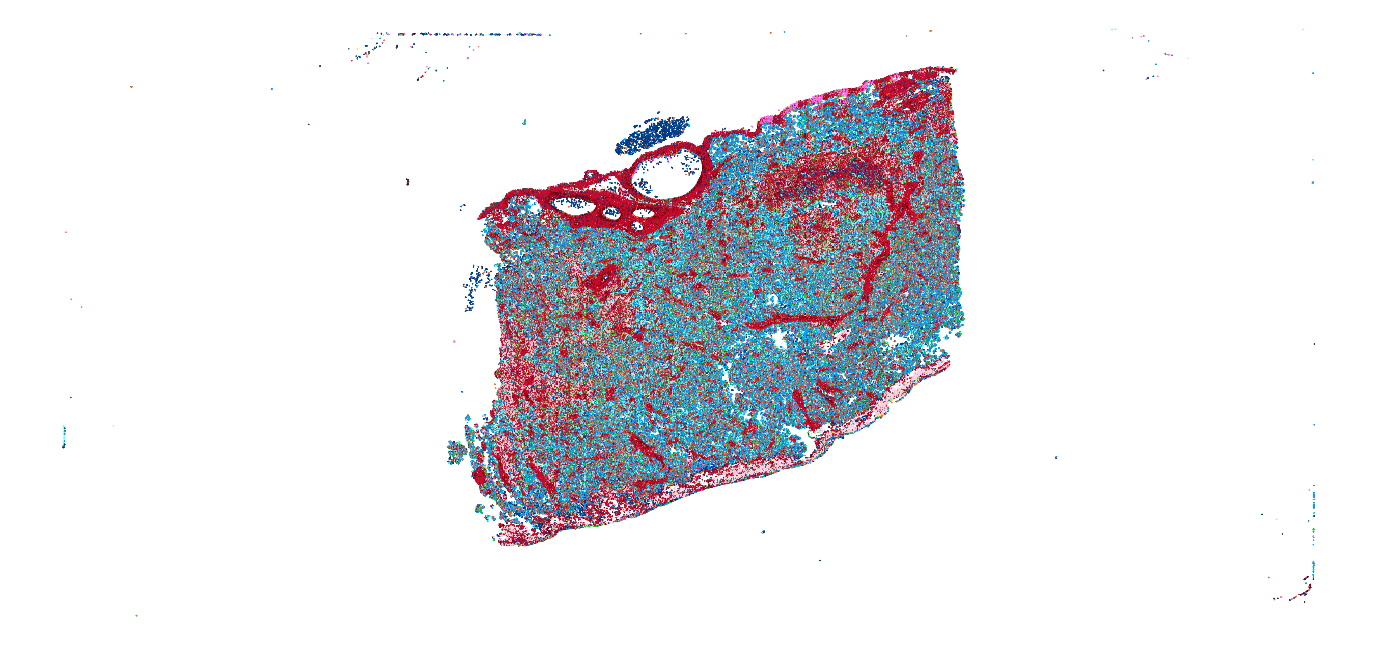

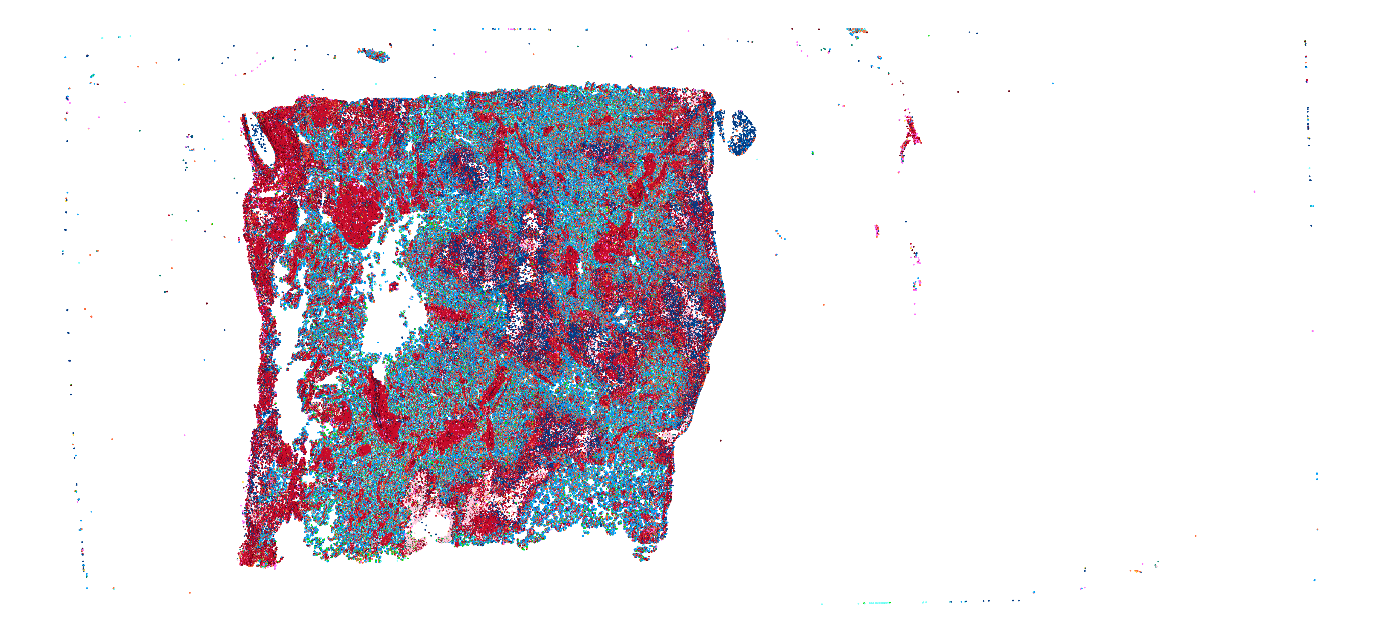

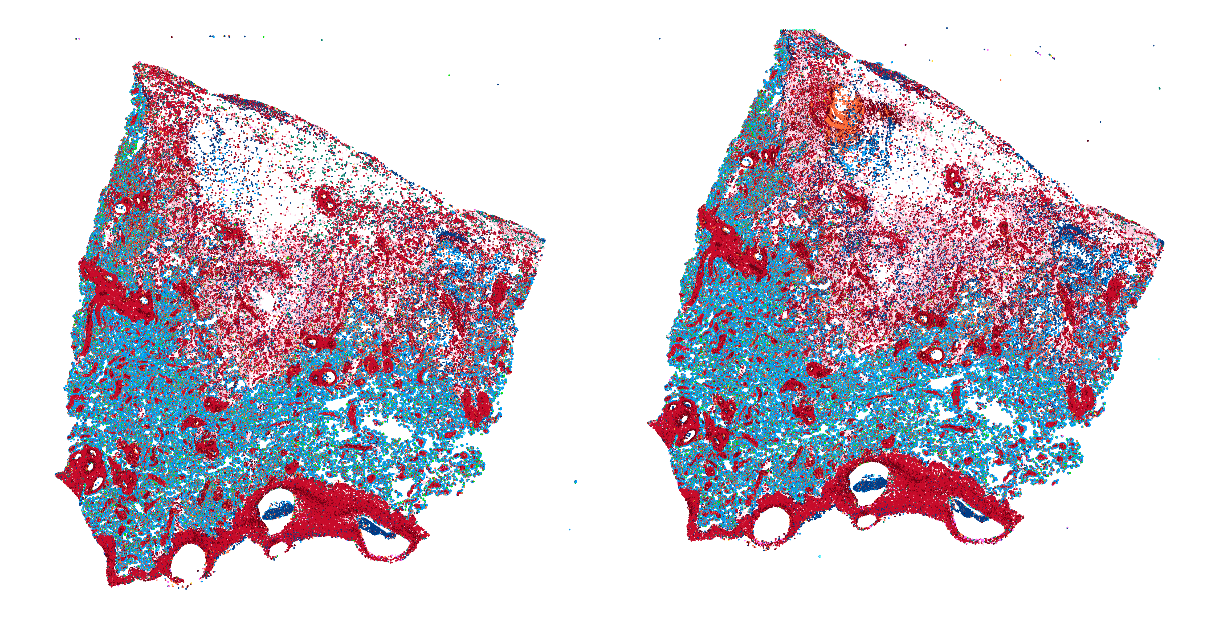

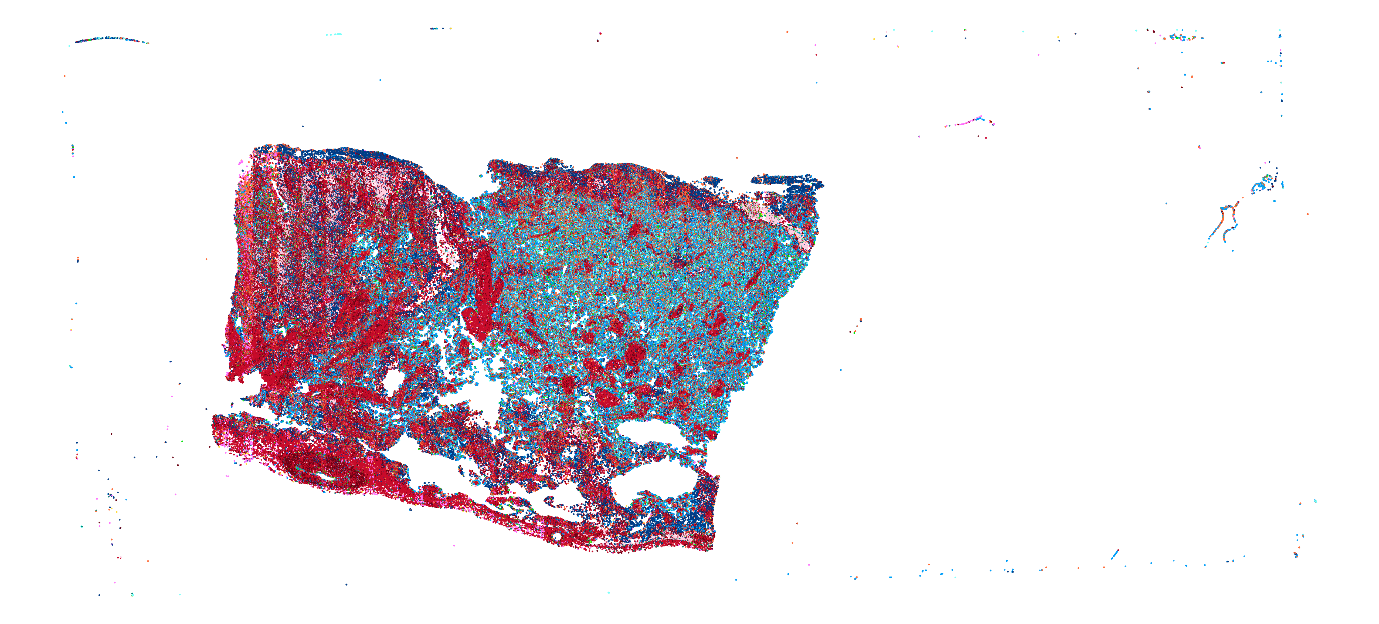

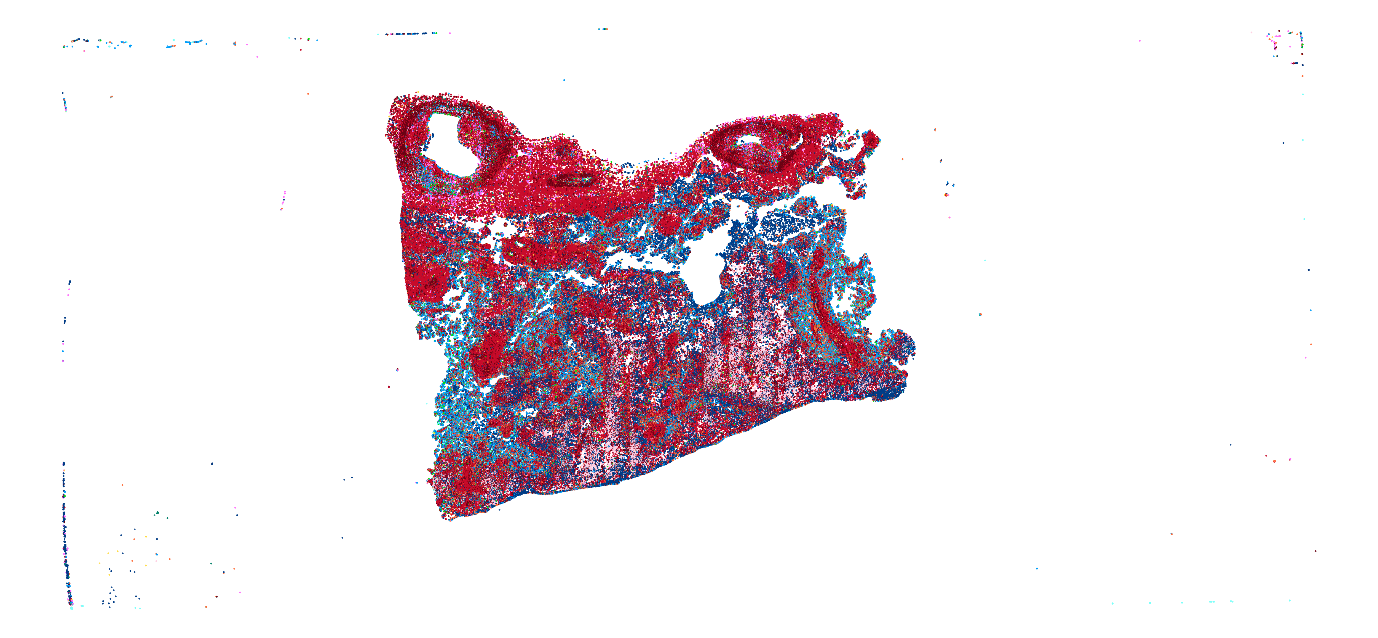

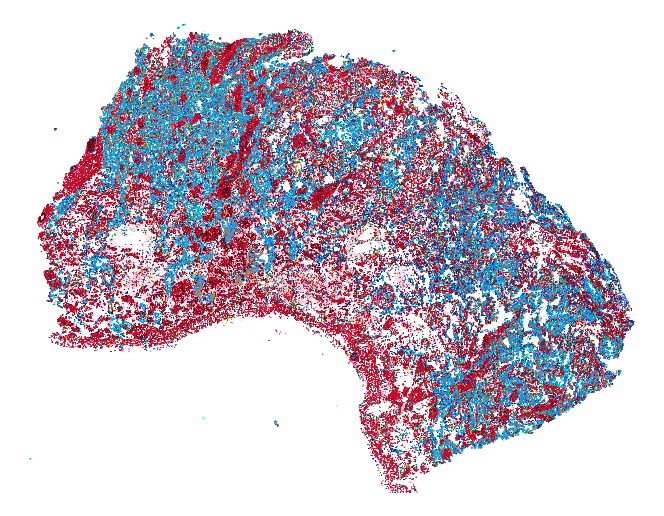

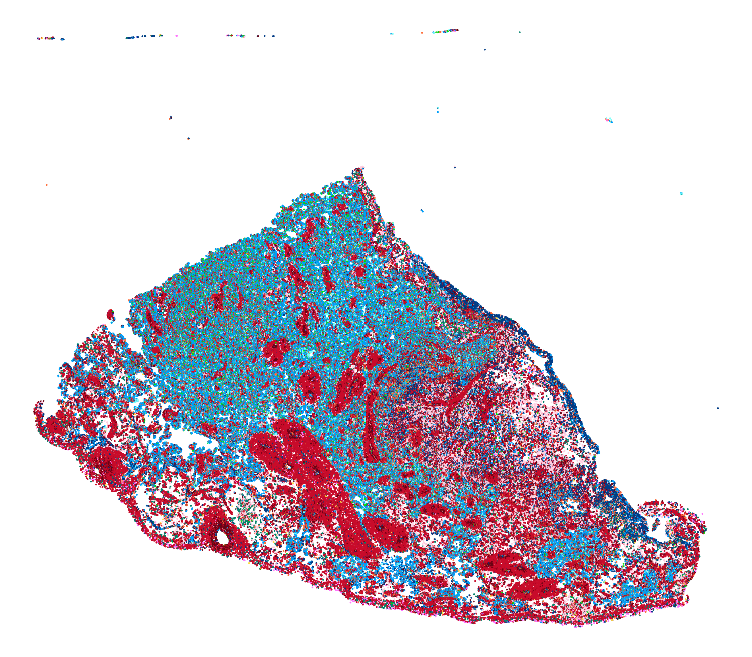

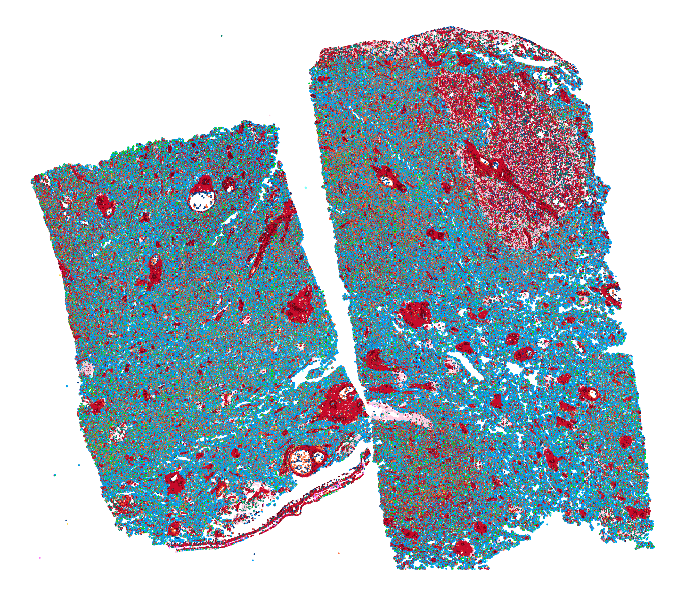

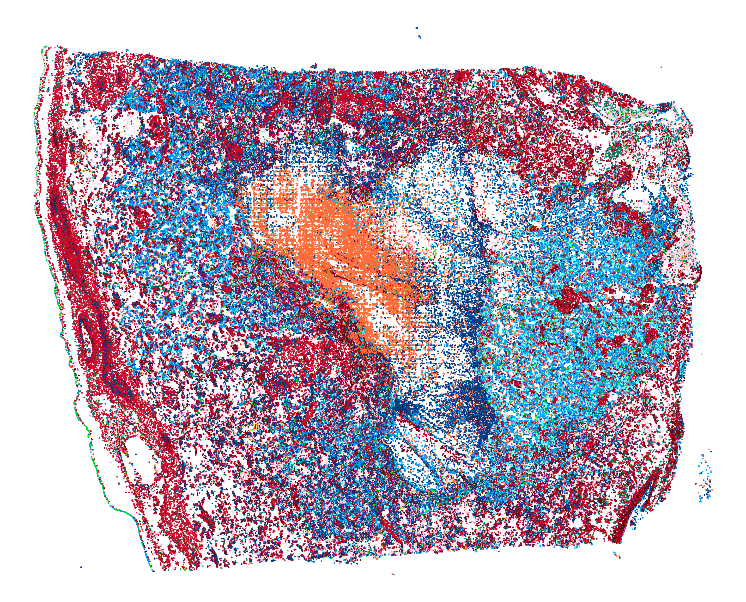

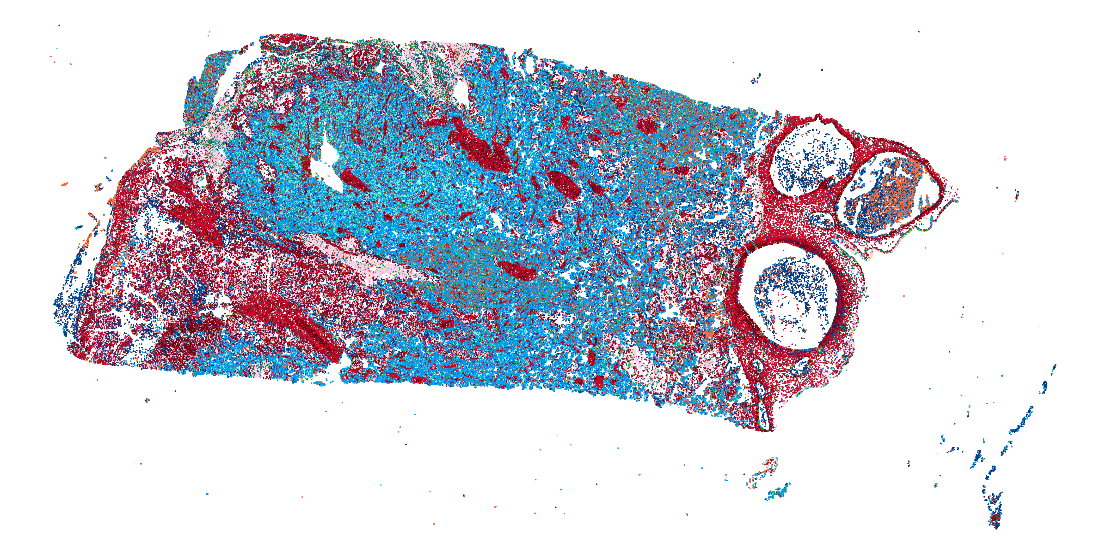

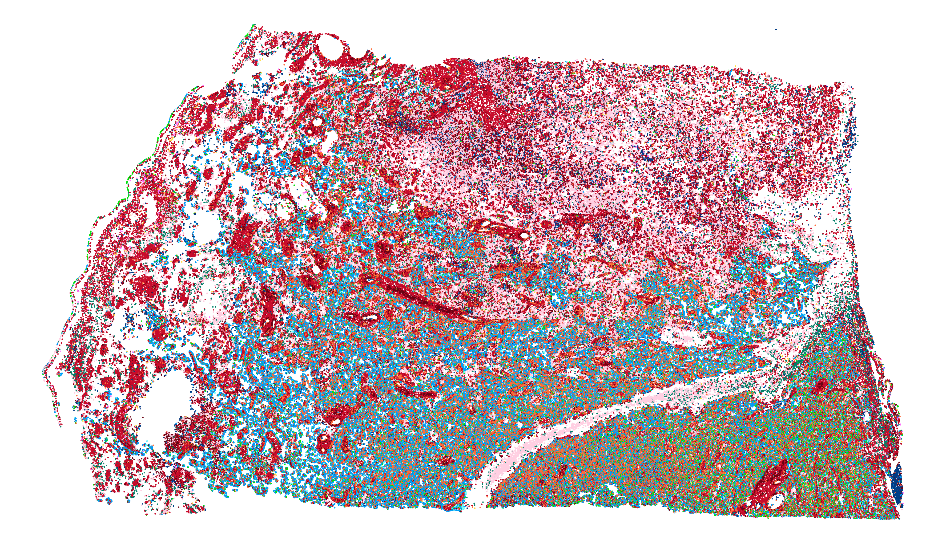

**Supplementary Fig. 4: Visualisation of nuclei and cell predictions across 12 WSIs of term placenta parenchyma with clinically significant placental infarction.**

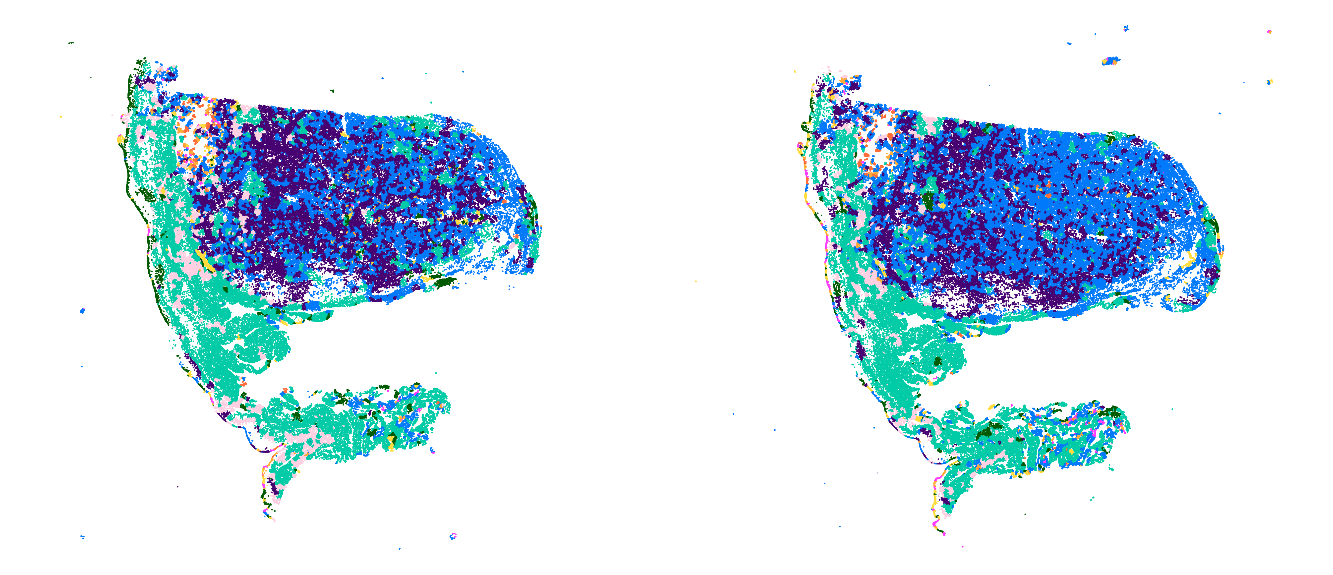

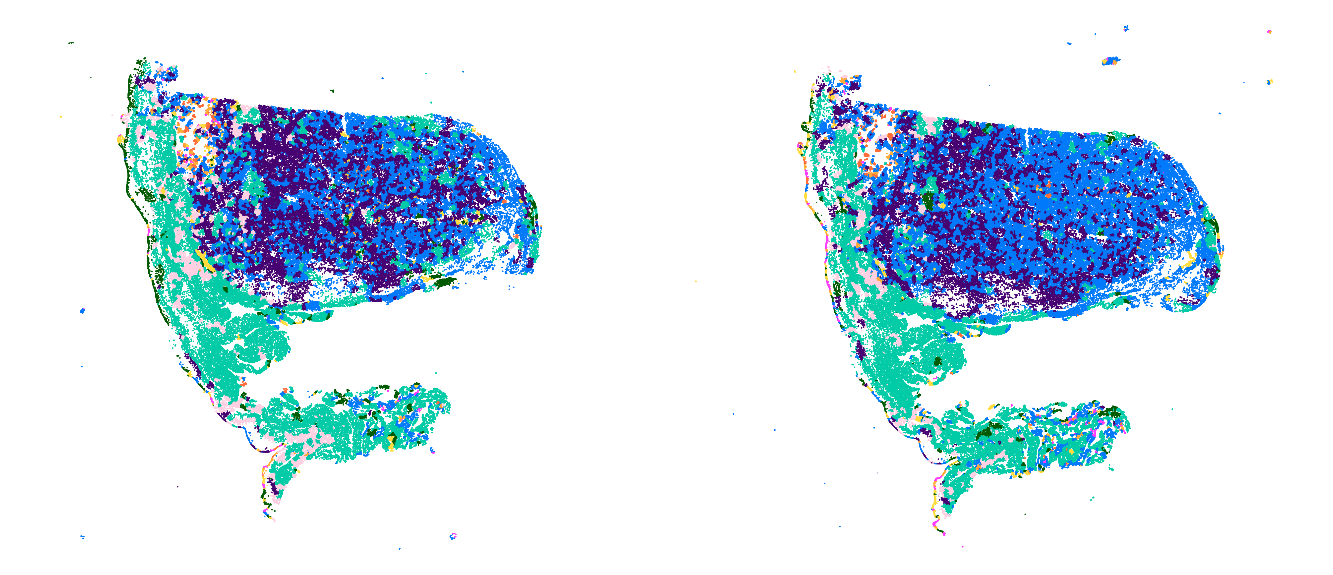

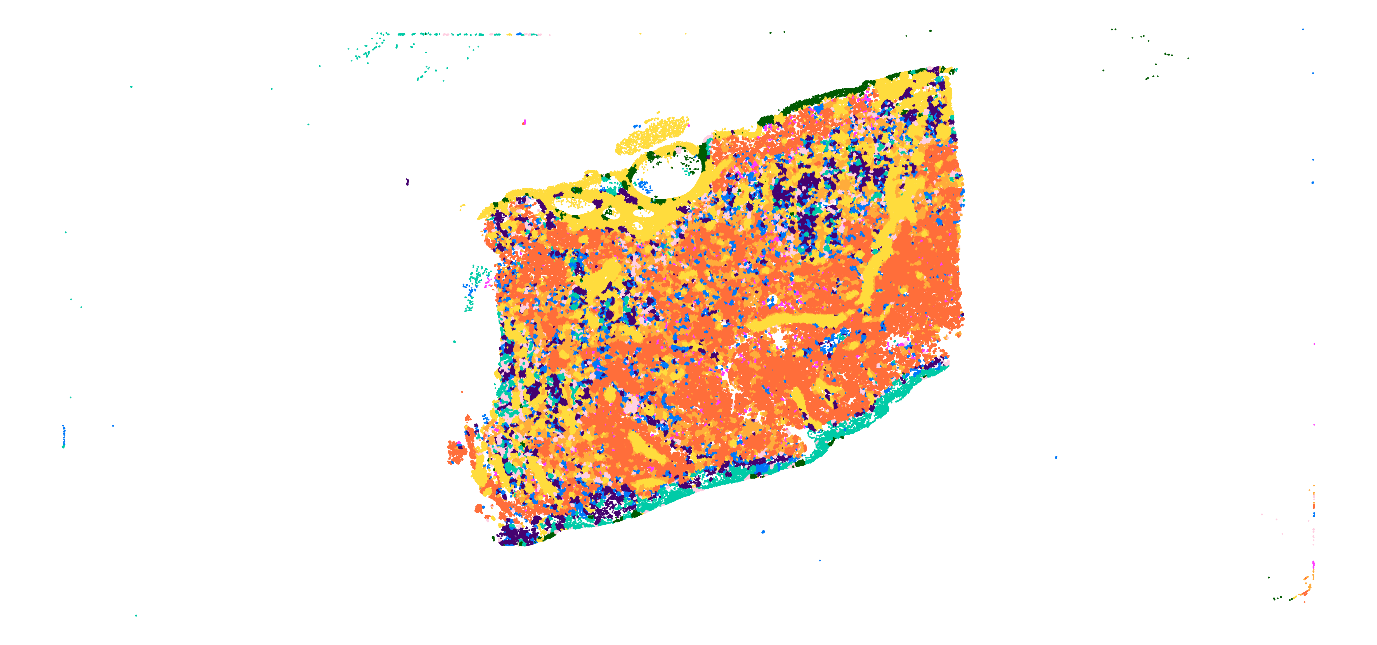

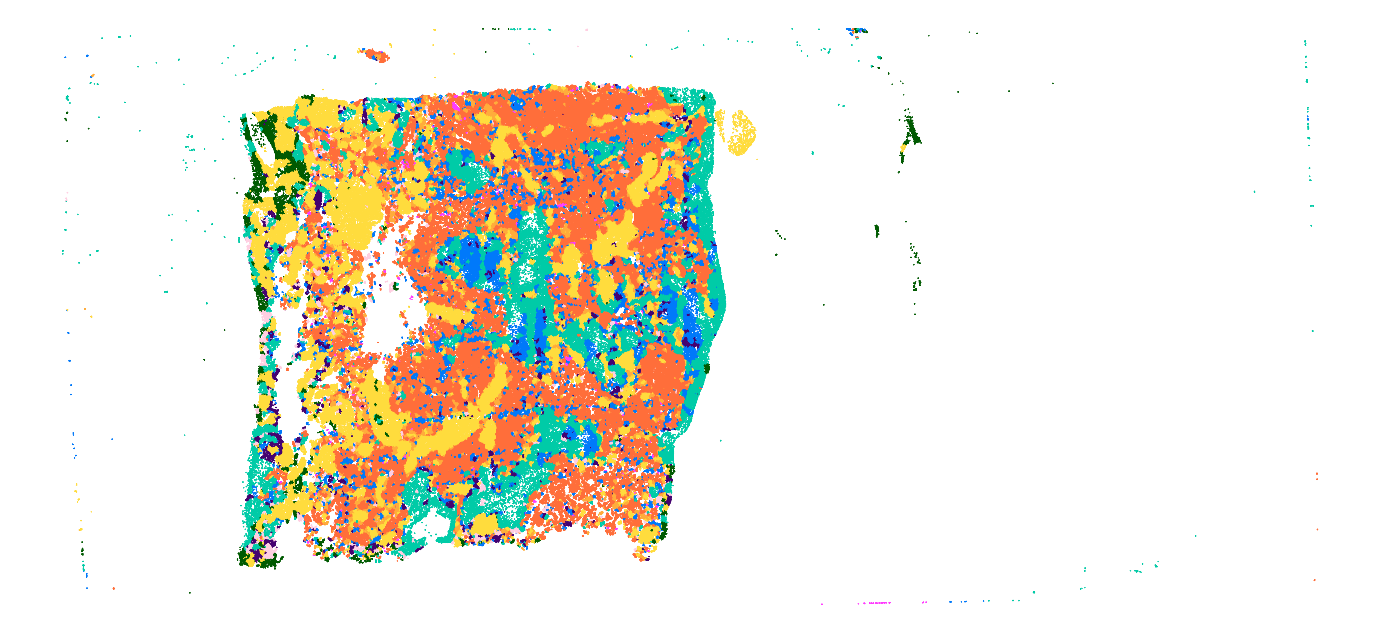

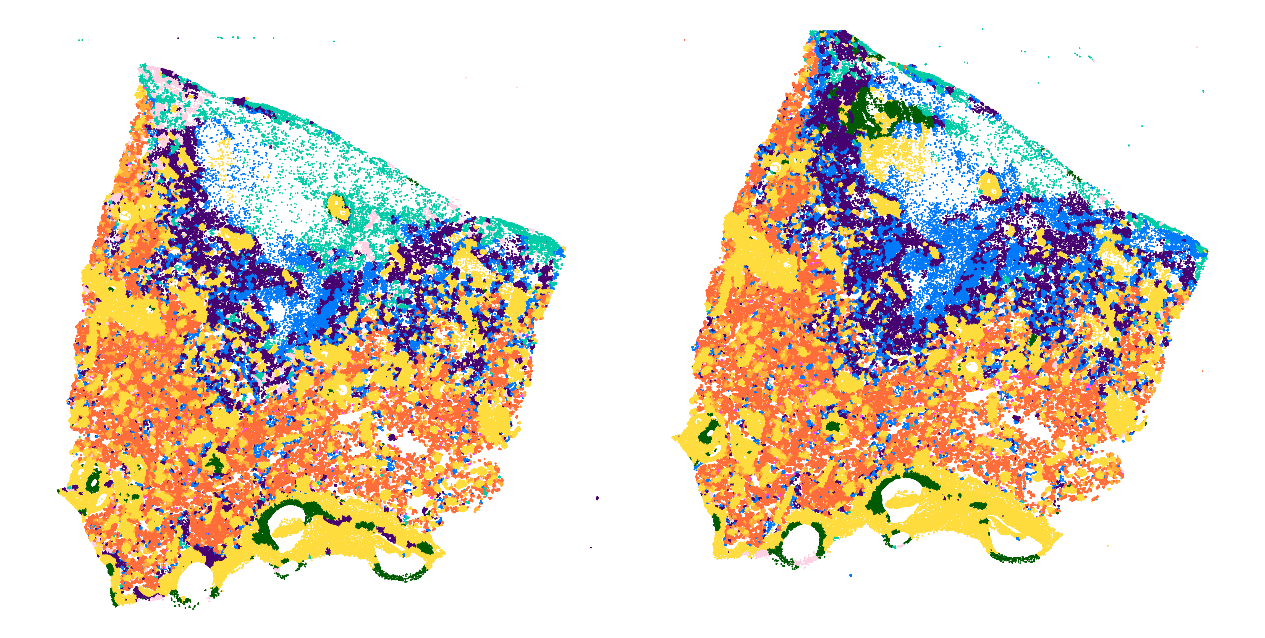

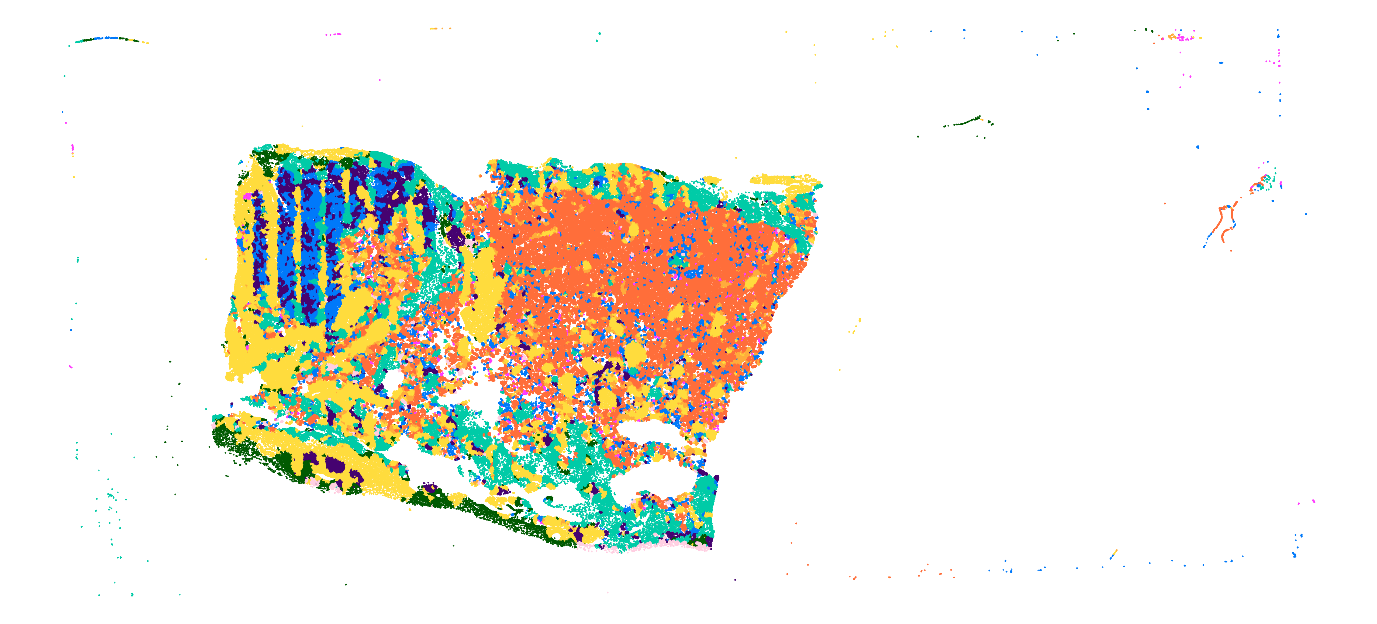

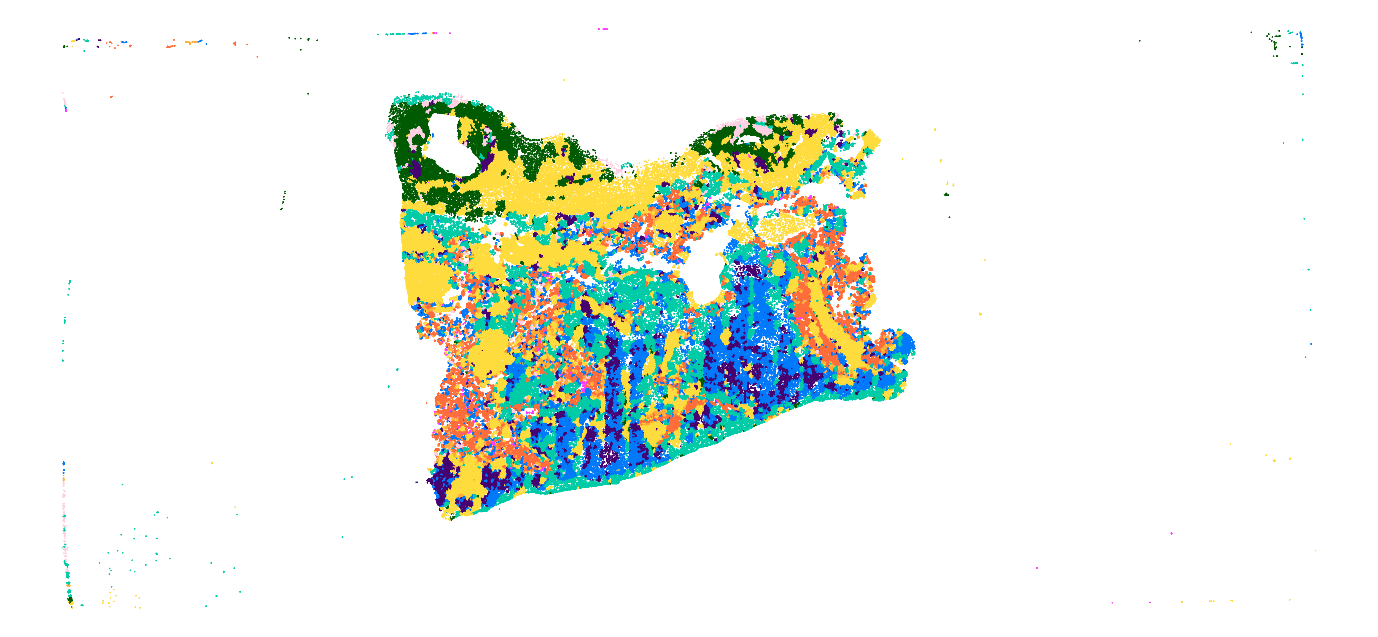

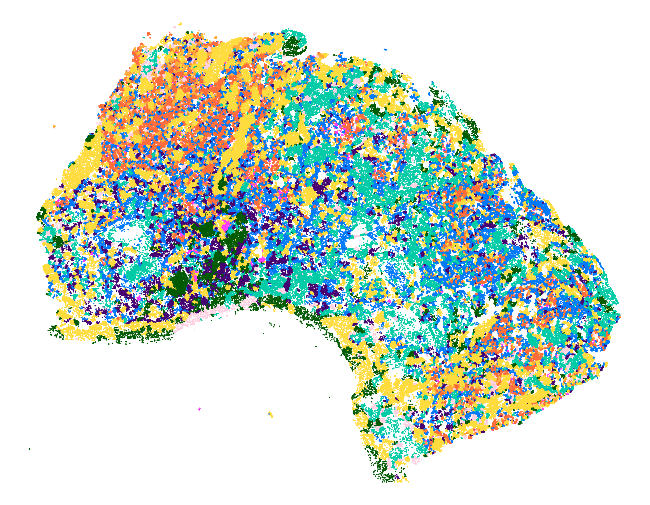

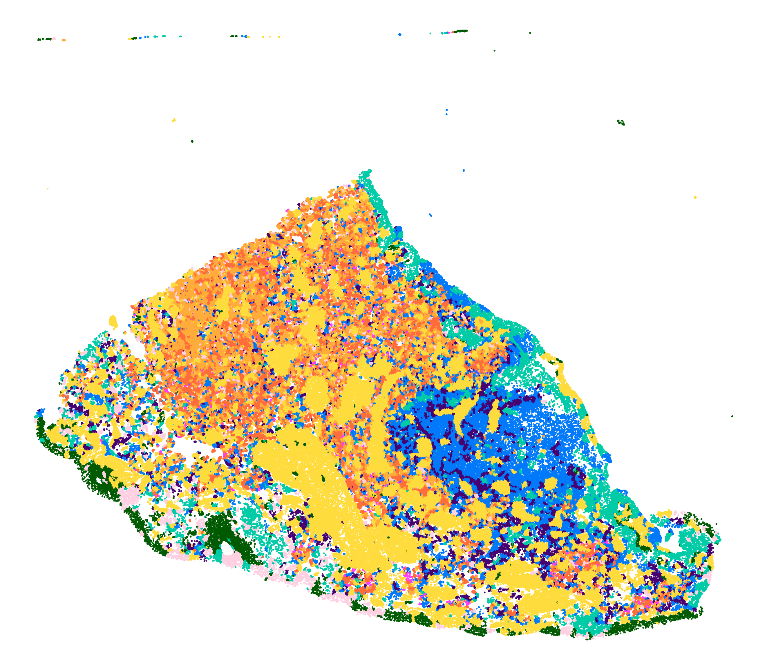

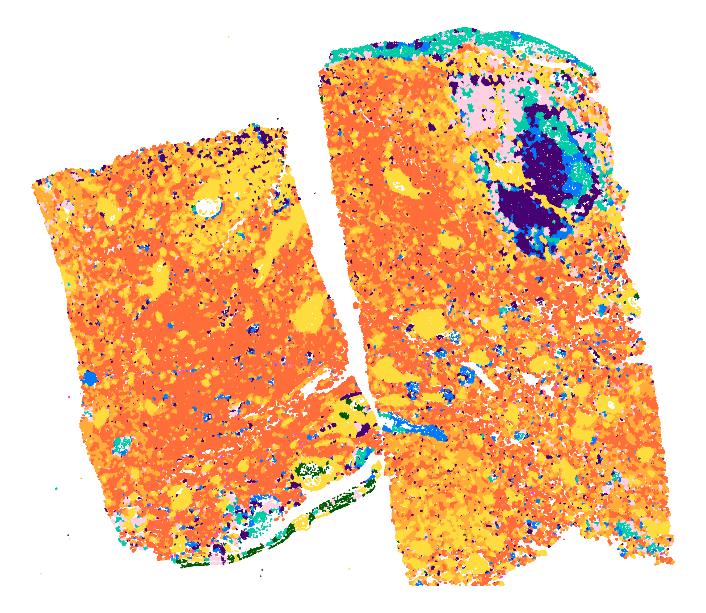

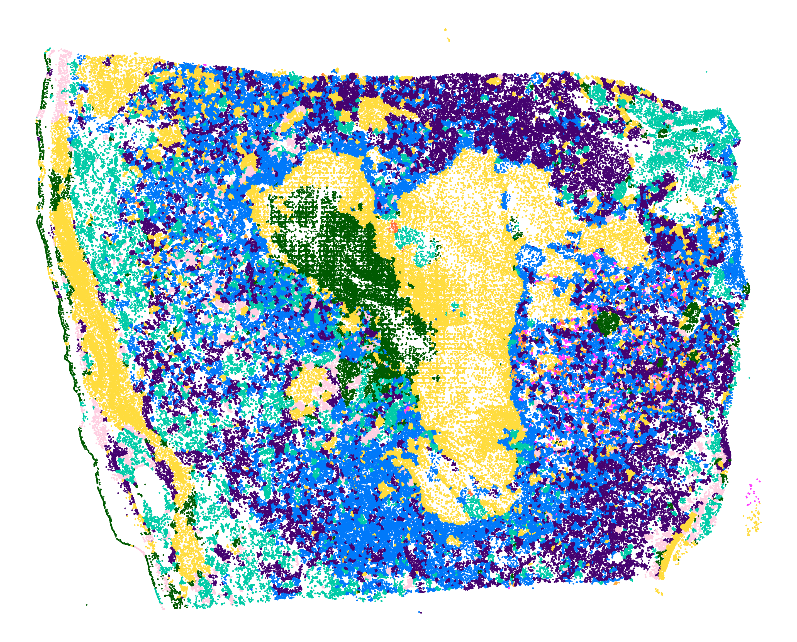

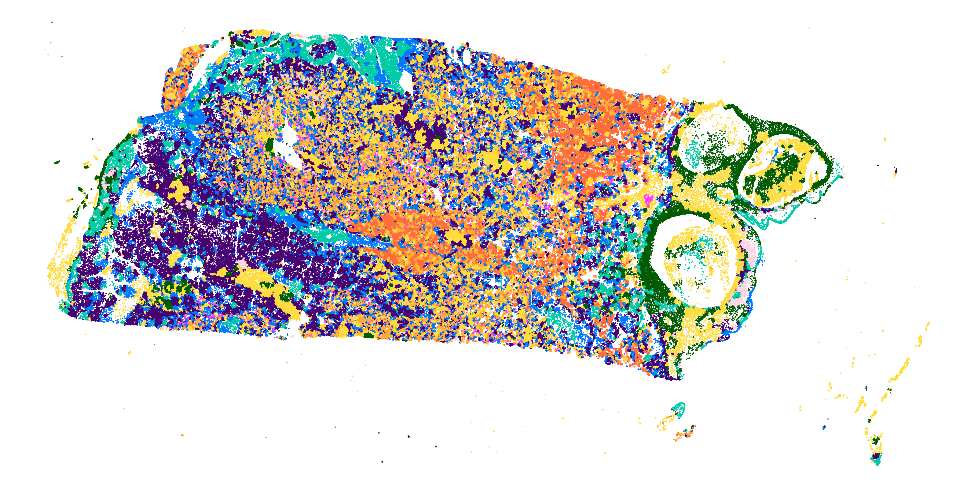

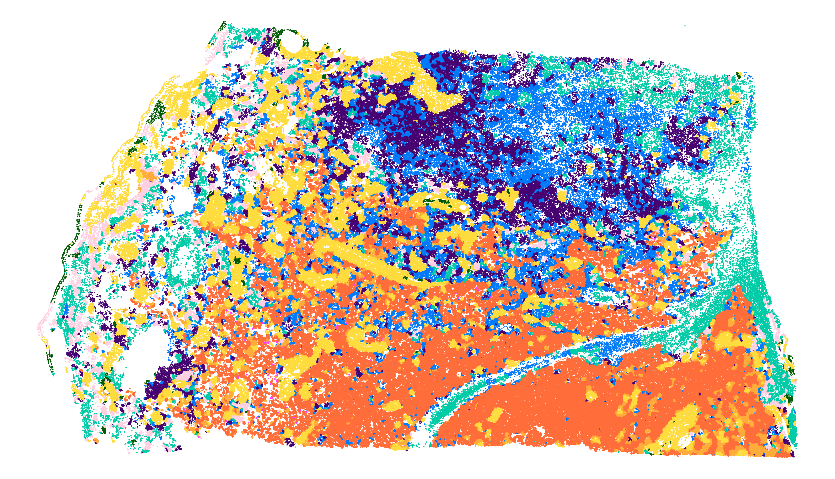

**Supplementary Fig. 5: Visualisation of nuclei and cell predictions across 12 WSIs of term placenta parenchyma with clinically significant placental infarction.**

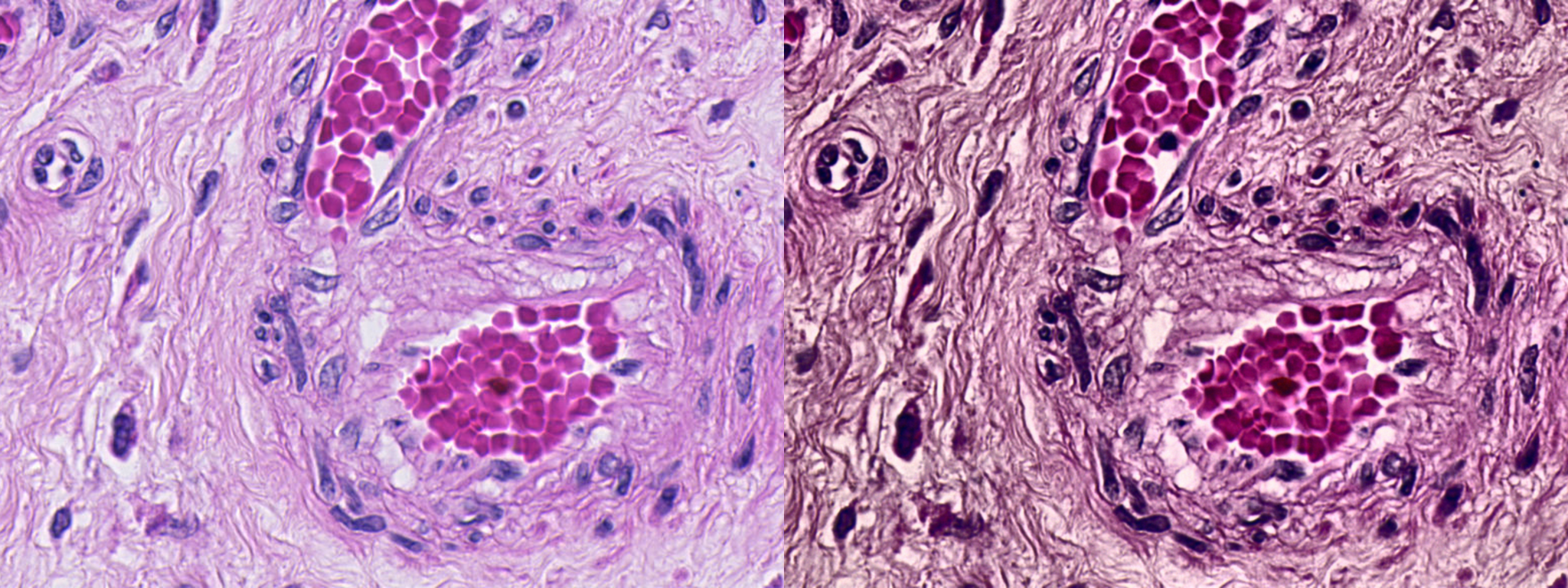

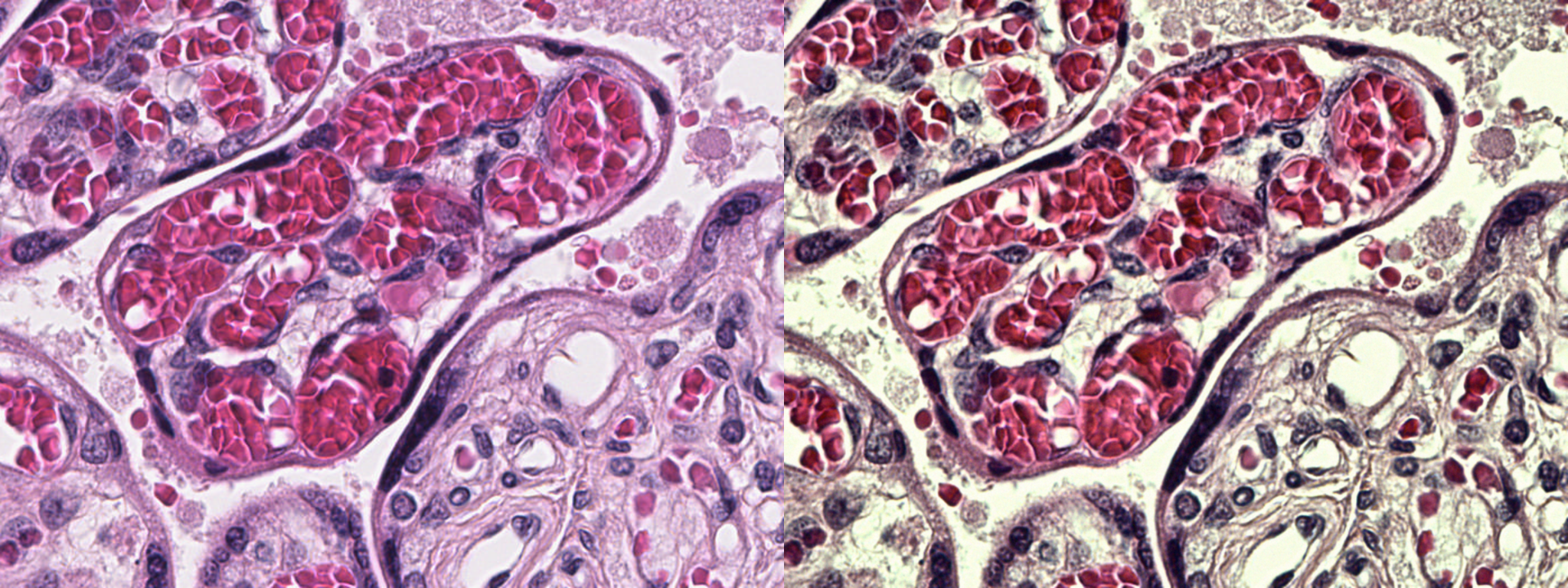

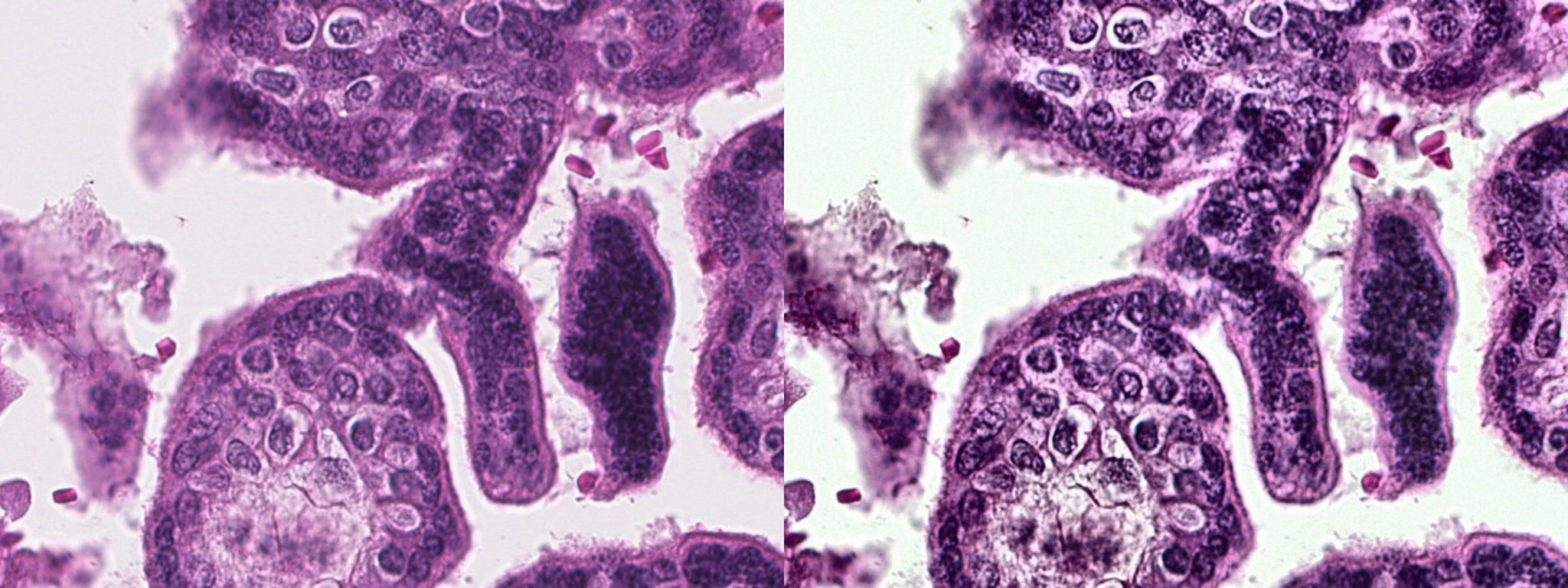

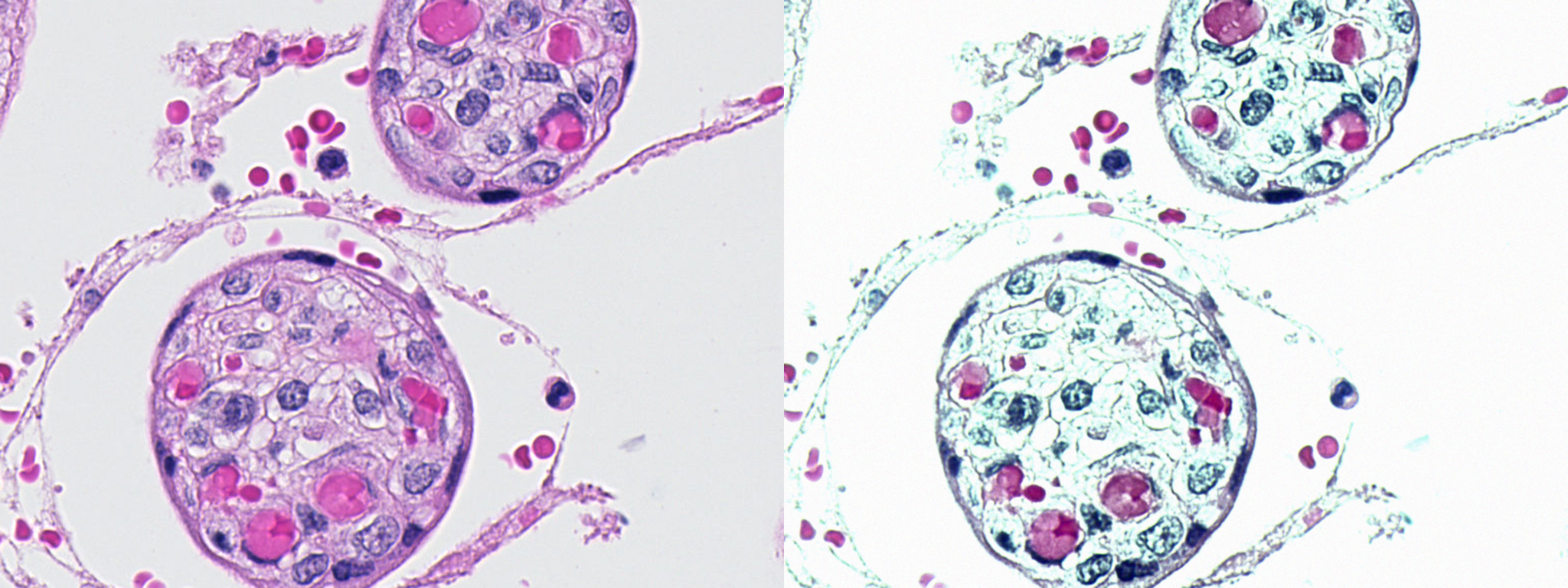

**Supplementary Fig. 6: Examples of H&E stain augmentation applied to nuclei localisation training data.** For each pair of images, the original histology image is on the left and the stain augmented histology image is on right.

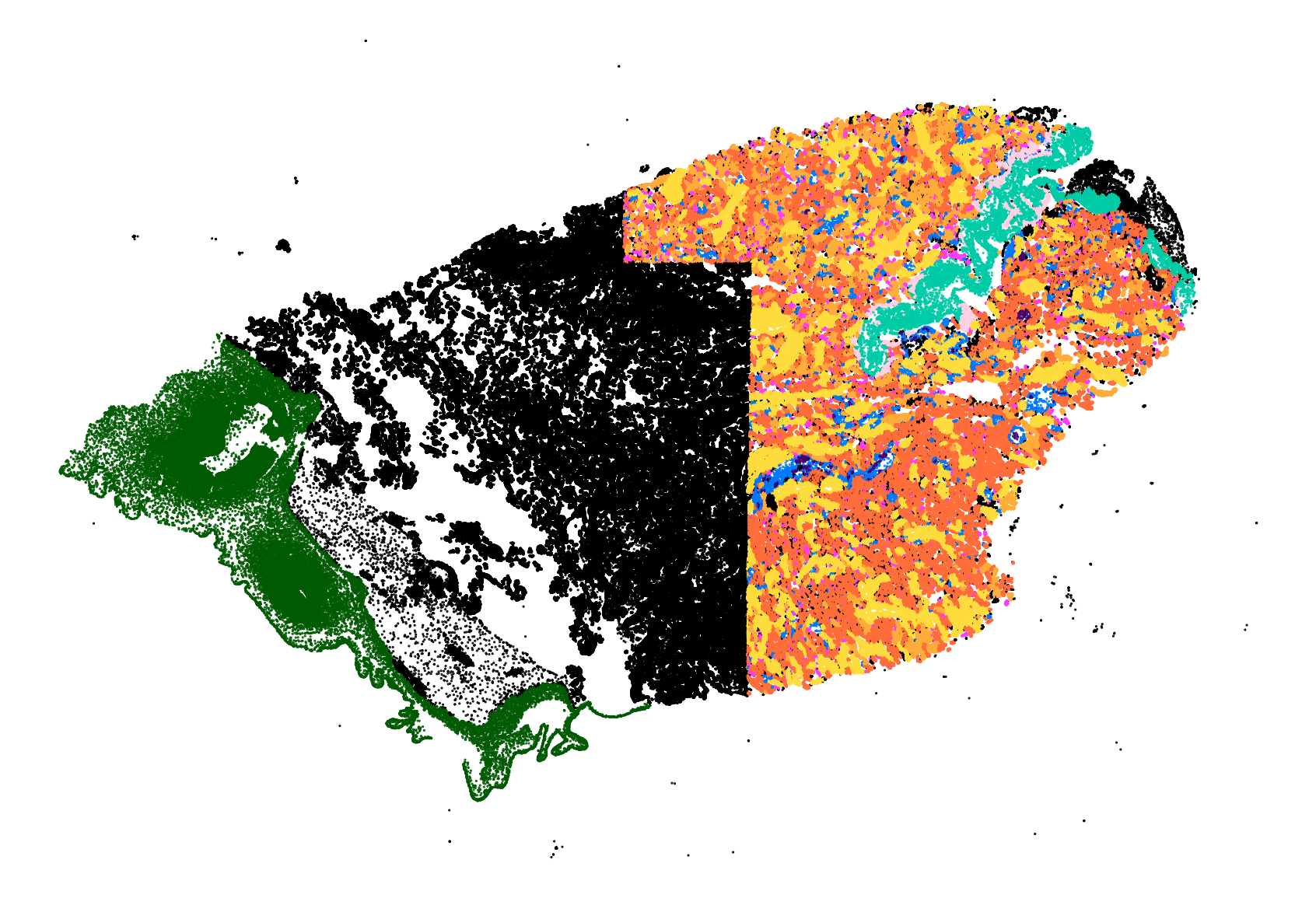

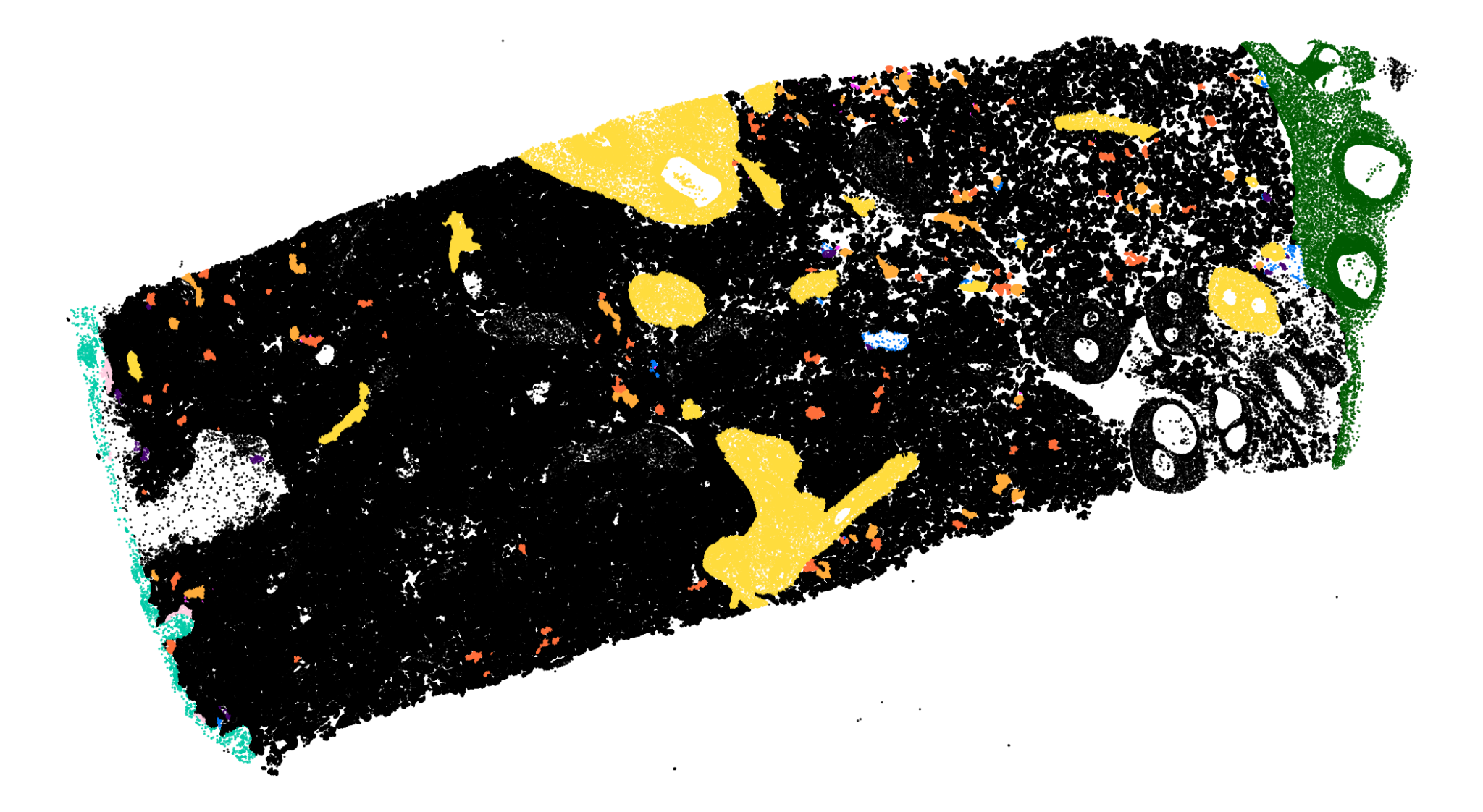

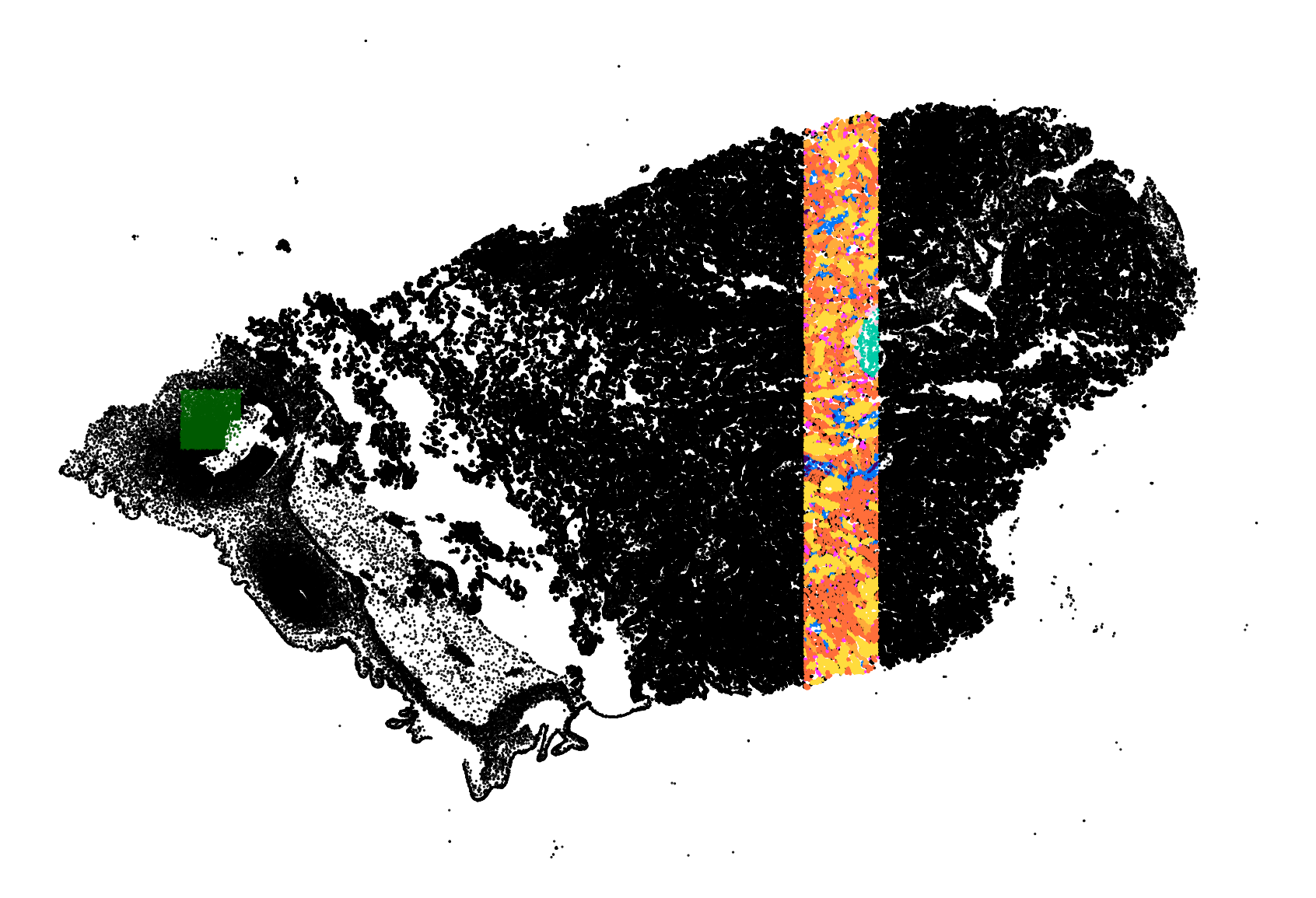

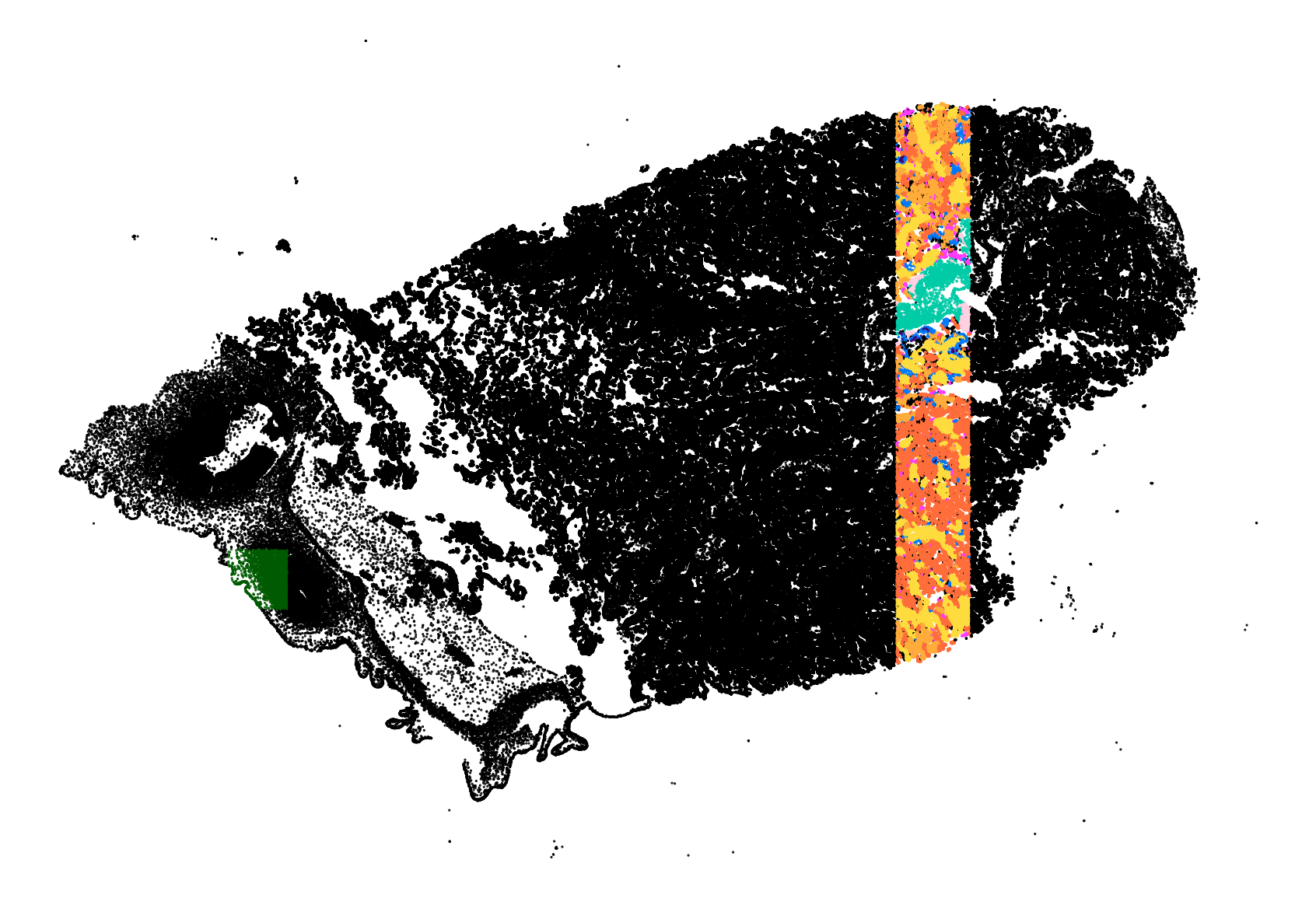

**Supplementary Fig. 7: Tissue node classification labelled data and validation and test split regions of WSI.** All labelled ground truth tissue node classification data across two WSIs (top) and validation (bottom left) and test (bottom right) split regions. Regions were chosen to contain a similar tissue type distribution to the training set and rest of the slide.

**Supplementary Tables 1-6**

**Table 1: The effect of stain augmentation on model generalisability.**

|  | **UoT1** | **UoT2** | **UoT3** | **HMC** |
| --- | --- | --- | --- | --- |
| **ND F1** | 0.89 | **0.896** | **0.829** | **0.868** |
| **ND_w/o_SA_ F1** | **0.899** | 0.882 | 0.812 | 0.843 |
| **CC AUC-PR** | 0.85 | 0.91 | 0.85 | **0.62** |
| **CC_w/o_SA_ AUC-PR** | **0.87** | **0.93** | **0.91** | 0.43 |

To examine the effect of our stain augmentations on the stain invariance of the nuclei detection (**ND**) and cell classification models (**CC**), we compare the performance of models trained with and without stain augmentations (**_w/o_SA_**) on datasets from different slides collected on different days and from two institutes (UoT and HMC). Models were trained with data from a set of slides at one institute (UoT1, UoT2, UoT3) and evaluated against unseen test data from this same institute and on unseen data from a second institute (HMC).

**Table 2: Term chorionic villus characteristics.**

| **Villus Type** | **Diameter (μm)** | **Trophoblast** | **Stroma** | **Vessels/capillary density** |
| --- | --- | --- | --- | --- |
| **Villus Sprout** | 60–100 | Syncytiotrophoblasts with a core of cytotrophoblasts | None or some reticular stroma | Unvascularised |
| **Terminal villi** | 60 | Vasculosyncytial membranes and syncitial knots | Scant fibrous stroma | Capillaries at more than 50% of the stromal area |
| **Mature Intermediate Villi** | 80–150 | Fully surrounded by syncytiotrophoblasts, half surrounded by cytotrophoblasts | Fibrous stroma | Capillaries at less than 50% of the stromal area |
| **Stem Villi** | 80–3000 | Thick layer, frequently replaced by fibrinoid | Fibrous stroma | Large muscularized arteries and veins, few capillaries |
| **Anchoring Villi** | 80-3000 | None or the same as stem villi | The same as stem villi | Unvascularised or the same as stem villi |

Characteristic and defining cellular and morphological traits of the term chorionic villus structures. Adapted from ^1,2^**.**

**Table 3: Patient characteristics for each placenta used for training.**

| **Institute** | **Models** | **Gestational Age** | **Relevant Clinical Data** |
| --- | --- | --- | --- |
| UoT | Nuclei and Cell | 30+5 | Acute chorioamnionitis, infarction |
| UoT | Nuclei and Cell | 30+5 | Acute chorioamnionitis, infarction |
| UoT | Nuclei and Cell | 40+2 | Umbilical cord knot, focal acute subchorionitis, intervillous thrombosis |
| UoT | Nuclei and Cell | 20+4 | Acute chorioamnionitis, inflammation of the umbilical cord |
| UoT | Nuclei and Cell | 42+0 | Placenta vallate, hypocoiled cord |
| HMC | Nuclei and Cell | 36+5 | Suspected placenta accreta |
| HMC | Nuclei and Cell | 41+5 | Suspected placenta accreta |
| UoT | Tissue | 42+0 | Placenta vallate, hypocoiled cord |
| HMC | Tissue | 40+0 | Suspected placenta accreta |

Patient characteristics and relevant clinical data for each placenta histology slide used for training. Slides used to train the nuclei and cell models have a range of pathologies and gestational ages, slides used to train the tissue model were from healthy term placentas.

**Table 4: Patient characteristics for each placenta used for inference.**

| **Institute** | **Gestational Age** | **Relevant Clinical Data** |
| --- | --- | --- |
| UoT | 42+0 | Placenta vallate, hypocoiled cord |
| HMC | 40+0 | Suspected placenta accreta |
| HMC | 36+5 | Suspected placenta accreta |
| HMC | 40+1 | Suspected placenta accreta |
| HMC | 41+5 | Suspected placenta accreta |
| UoT | 40+5 | Infarction, intervillous thrombosis |
| HMC | 40+4 | Infarction, umbilical vessel inflammation |
| HMC | 40+0 | Infarction, intervillous thrombosis |
| HMC | 40+1 | Infarction, perivillous fibrin |
| HMC | 41+1 | Infarction, perivillous fibrin, accreta |
| HMC | 40+4 | Infarction, chorioamnionitis |
| HMC | 40+6 | Infarction, perivillous fibrin, avascular villi, chronic villitis, accreta |
| HMC | 41+1 | Infarction, perivillous fibrin, intervillous thrombosis, avascular villi, chronic villitis, accreta |

Patient characteristics and relevant clinical data for each placenta used for inference. Slides from the first five placentas correspond to the healthy group presented in the results and slides from the last eight placentas correspond to the group with clinically significant placental infarction.

**Table 5: Training augmentations for nuclei localisation and cell classification.**

| **Augmentation** | **Probability** | **Additional Parameters** |
| --- | --- | --- |
| Flip | 0.5 | - |
| RandomRotate90 | 0.5 | - |
| StainAugment | 0.9 | variance=0.4 |
| CLAHE | 0.8 | clip limit=3.0, tile_grid_size=(8, 8) |
| GaussNoise | 0.8 | var_limit=(10.0, 200.0) |
| Blur | 0.8 | blur_limit=5 |

Augmentations are applied sequentially using the Albumentations library^3^ except for StainAugment. StainAugment is a custom stain deconvolution method which varies H&E stain intensities using one of 8 randomly chosen RGB to H&E colour matrices^4^. Colour matrices are drawn from the original literature and derived from our own slides.

**Table 6: Datasets and Dataset Splits.**

| **Nuclei Detection** | | | |
| --- | --- | --- | --- |
|  | Train | Validation | Test |
| Nuclei | 11755 (70%) | 2374 (14%) | 2754 (16%) |
| Patches | 176 (70%) | 38 (15%) | 38 (15%) |
| **Cell Classification** | | | |
| Syncytiotrophoblast | 2750 (70%) | 583 (15%) | 548 (15%) |
| Cytotrophoblast | 887 (70%) | 184 (15%) | 175 (15%) |
| Syncytial Knot | 845 (72%) | 146 (12%) | 179 (16%) |
| Extravillus Trophoblast | 968 (70%) | 178 (14%) | 169 (16%) |
| Fibroblast | 2260 (70%) | 423 (14%) | 427 (16%) |
| Hofbauer Cell | 194 (72%) | 35 (13%) | 40 (15%) |
| Vascular Endothelial | 1292 (70%) | 252 (14%) | 248 (16%) |
| Vascular Myocyte | 1492 (70%) | 323 (15%) | 329 (15%) |
| Mesenchymal Cell | 218 (67%) | 58 (18%) | 47 (15%) |
| Maternal Decidua | 151 (68%) | 37 (17%) | 35 (16%) |
| Leukocyte | 732 (69%) | 142 (13%) | 184 (17%) |
| **Tissue Classification** | | | |
| Villus Sprout | 15545 (50%) | 9066 (29%) | 6441 (21%) |
| Terminal Villi | 159464 (50%) | 84052 (27%) | 72402 (23%) |
| Mature Intermediate Villi | 80954 (51%) | 39296 (25%) | 37332 (24%) |
| Stem Villi | 132138 (55%) | 58658 (25%) | 48568 (20%) |
| Anchoring Villi | 5769 (64%) | 1634 (18%) | 1654 (18%) |
| Chorionic Plate | 46584 (84%) | 4942 (9%) | 3937 (7%) |
| Basal Plate and Septum | 13329 (75%) | 1029 (6%) | 3359 (19%) |
| Fibrin | 14195 (54%) | 7047 (27%) | 5010 (19%) |
| Avascular Villi | 891 (49%) | 526 (29%) | 392 (22%) |

Number of nuclei, cells, and tissue nodes used for training, validation and test sets of respective models followed by, in brackets, the proportion of data for that class across splits.
