## Supplementary File 1 for "HAPPY: A deep learning pipeline for mapping cell-to-tissue graphs across placenta histology whole slide images"

Placenta Human Validation SOP

### Purpose

To assess the levels of agreement/disagreement between the author as a self-taught but untrained annotator (*original annotator*) to those of practising perinatal pathologists on H&E stained placenta histology substructures. Substructures of interest are split into two categories: tissues and other large structures (i.e. terminal villus or areas of fibrin) and cells (i.e. syncytiotrophoblast).

Each participating pathologist will be anonymous and will not be able to see the choices or interactions of other pathologists throughout the validation process. No data which can be linked to any one individual will be shared.

Ultimately, the original annotator’s annotations will be used to train and validate a deep learning approach to modelling placenta substructures. Indirectly, the inputs gathered from perinatal pathologists will form part of this validation for the deep learning models.

### Setup/Design

##### **Whole Slide Image and Patient Selection**

The whole slide image chosen for validation is an H&E stained parenchyma section of a term placenta from an uncomplicated pregnancy, without adverse maternal or neonatal outcomes. Pathologist’s notes from the delivery site write that this section is histologically normal and “inflammation free… with villi corresponding to gestational age”.

##### **Substructures of Interest**

| **Tissue Structures** | **Cells** |
| --- | --- |
| Terminal Villus | Syncytiotrophoblast |
| Mature Intermediary Villus | Cytotrophoblast |
| Immature Intermediary Villus | Fibroblast |
| Stem Villus | Hofbauer Cell |
| Anchoring Villus | Vascular Endothelial Cell |
| Mesenchymal Villus | Vascular Myocyte |
| Villus Sprout | Maternal Decidual Cell |
| Chorion/Amnion | Extra Villus Trophoblast |
| Avascular Villus | Leukocyte |
| Basal Plate/Septa | Mesenchymal Cell |
| Fibrin |  |
| Inflammatory Response |  |

Substructure References:

- <https://link.springer.com/chapter/10.1007/978-3-030-11425-1_36>
- <https://www.proteinatlas.org/learn/dictionary/normal/placenta>
- <https://www.cambridge.org/core/books/placental-and-gestational-pathology/normal-development/11AA85DC1FF68EDA1165A2B46102F1D2>
- <https://onlinelibrary.wiley.com/doi/10.1111/apm.12858>

##### **Software Used**

Original annotations and images were made using QuPath v3.1 (<https://qupath.github.io/>), images and collected data are stored securely in the Google Cloud Platform (<https://cloud.google.com/>), and the human validation interface is built using LabelBox (<https://labelbox.com/>).

##### **Tissue Validation Process**

Each pathologist is invited to join the LabelBox project. After signing in, they are presented with an image containing a centred single tissue structure or a cluster of assumed same-type tissue structures. They have several options per image, they can:

- view a polygon highlighting the area of interest (see image)
- set the tissue structure label and submit it
- say the tissue structure is unclear from the image and skip it
- say the tissue structure isn’t in the list of options
- add a text comment to the image

Once they have submitted their label choice, they are presented with the next image in the queue. Images are queued such that all pathologists are shown the same images in the same order and the queue contains an even, but random, distribution of each tissue class originally annotated on the term sample. The exact number of each tissue class is not disclosed to avoid biasing the pathologists.

A tutorial detailing the validation process may be found here:[LabelBox Tissue Validation Guide](https://docs.google.com/document/d/1IZYIaipnW-1-73sX10k70ZIj978NirZOsmYqWwmlDPE/edit?usp=sharing)

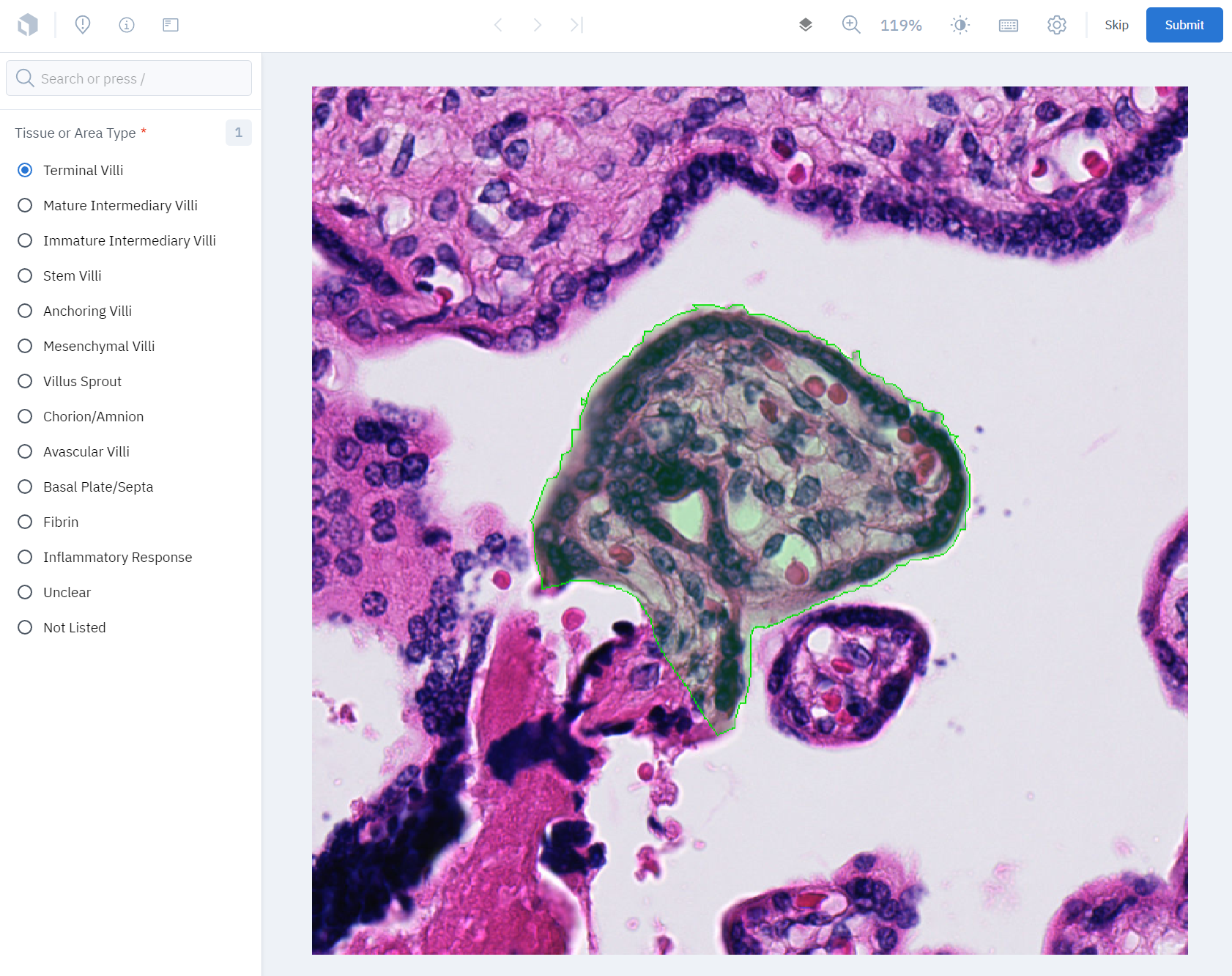

##### **Tissue Image Generation**

1. Polygons around structures were drawn using QuPath’s Pixel Classifier and then corrected by the original annotator.
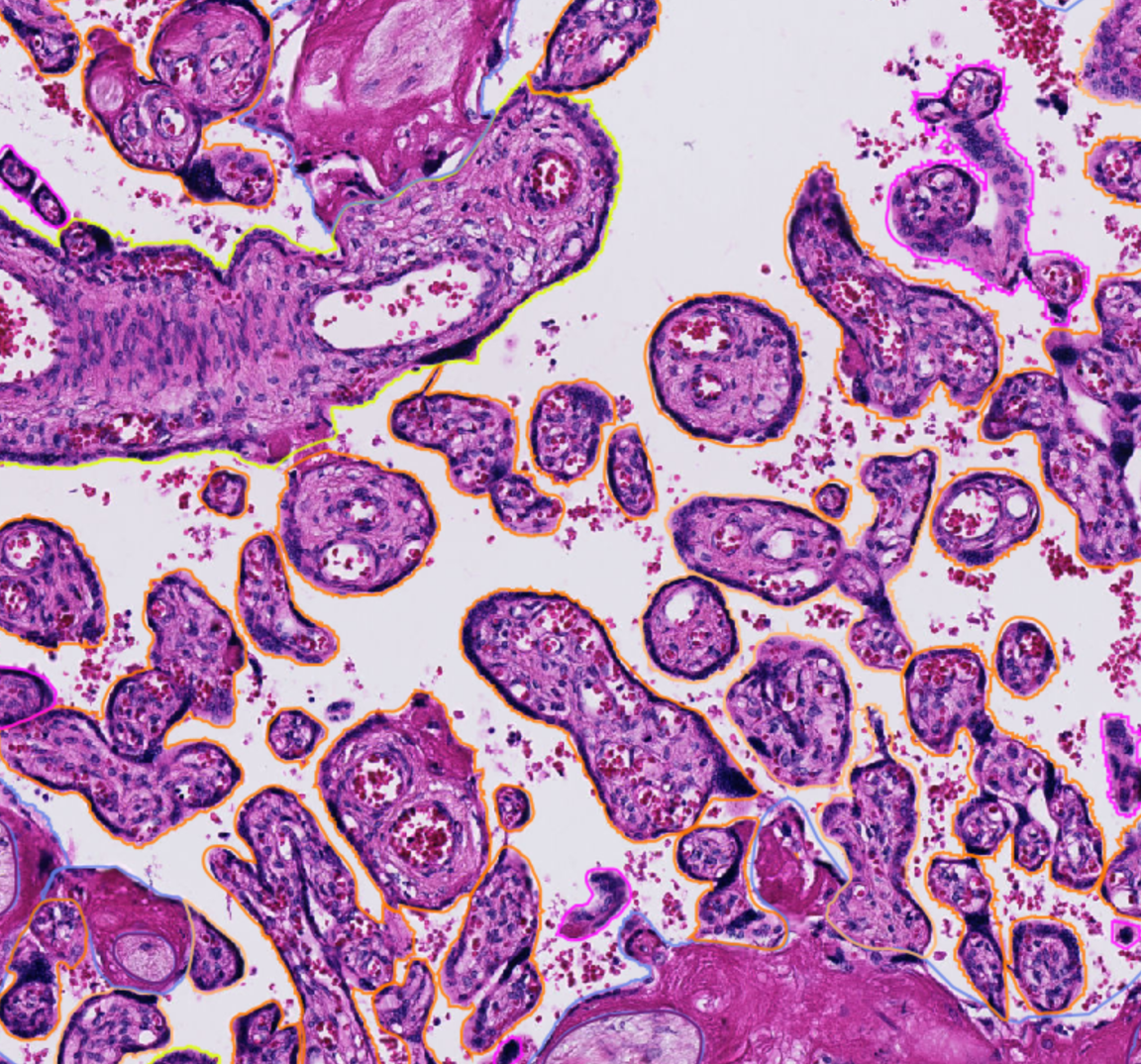

2. Tissue structure classes were assigned by the original annotator.
3. A downsampled image of the tissue boundaries was extracted so that each image contained a centred tissue or a cluster of same-type tissues.
4. For the larger structures such as the chorion, multiple images were instead extracted from smaller regions.
5. The corresponding coordinates of the polygons were saved in a json file.
6. Duplicate images with overlaid polygons were generated.
7. Images and polygon images were uploaded to Google Cloud and linked to LabelBox.

##### **Data Collected for Validation**

- Chosen tissue structure label
- Time spent on each image
- Number of images submitted
- Number of images submitted as ‘unclear’ or ‘not listed’
- Optional comments submitted by pathologists
  - Comments will remain anonymous and will not be shared or quoted publicly

##### **Downstream Analysis**

The data collected from pathologist validators will predominantly be used to assess the untrained original annotator. The annotator’s performance will be judged by the amount of agreement between their original annotations and the pathologists’ label choices. The agreement will be quantified using Cohen’s Kappa across all labelled data and also individually for each tissue type.

Agreement between pathologists will be quantified using the same metrics to assess the relative difficulty of the task. Of particular interest is whether there are any consistently difficult tissue structures that have low agreement scores and whether there are structures that have common patterns of disagreement (i.e. is there high disagreement between terminal villi and mature intermediary villi, for example).

These patterns of disagreement, should they exist, will also be used as a benchmark against the performance of a deep learning model on the same task. For example, if there is a large disagreement between terminal villi and mature intermediary villi by pathologists, then we would expect a deep learning model to mistake the two tissues more commonly than it would terminal villi and stem villi.
