## Supplementary File 2 for "HAPPY: A deep learning pipeline for mapping cell-to-tissue graphs across placenta histology whole slide images"

**Placenta Tissue Validation Guide**

Thank you for taking the time to validate these placenta tissues!

Any questions or feedback, please

### **Instructions:**

- You will be shown images containing tissue structures.
- To see the original image without overlay, use the ‘Image overlay’ button and select ‘Layer 2’.
- Choose a tissue structure from the list on the left and ‘Submit’ the image.
- If you are unsure, change the tissue type to ‘Unclear’ and ‘Submit’.
- You may optionally leave a comment on the image.

Further instructions on how to **show the original image** (p1), **change the tissue type** (p2), **see placenta sample information** (p3), and **add a comment** (p4) are shown in the pages that follow.

If you would like a reference guide to **villus tissue structures** from the literature, see page 5.

**Show Original Image**

Click on the ‘Image overlay’ button in the top right and then select ‘layer 2’. You may use ‘layer 1’ to return the polygon overlay.

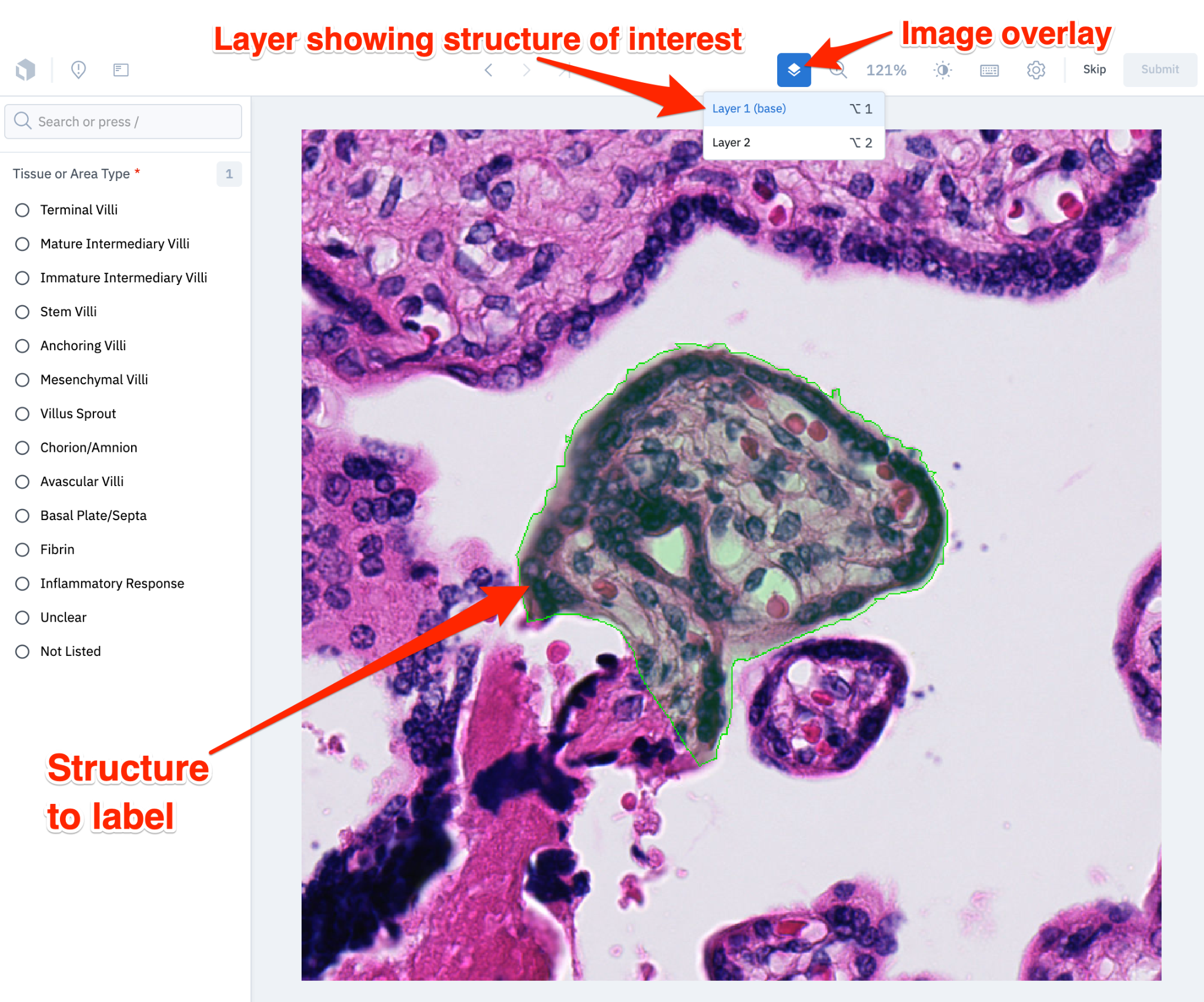

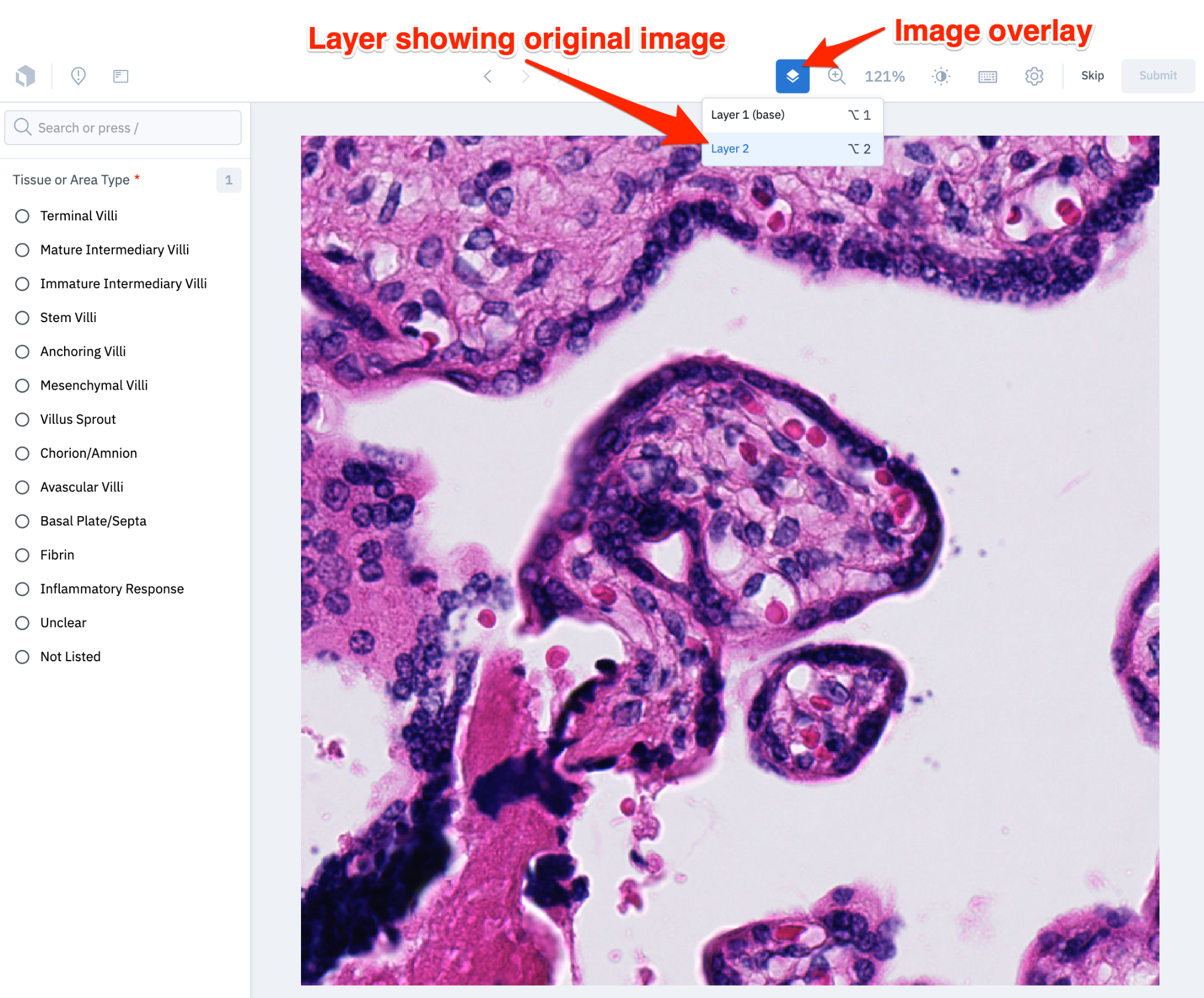

**Change the Tissue Type**

Choose an option from the list on the left and then press the ‘Submit’ button in the top right.

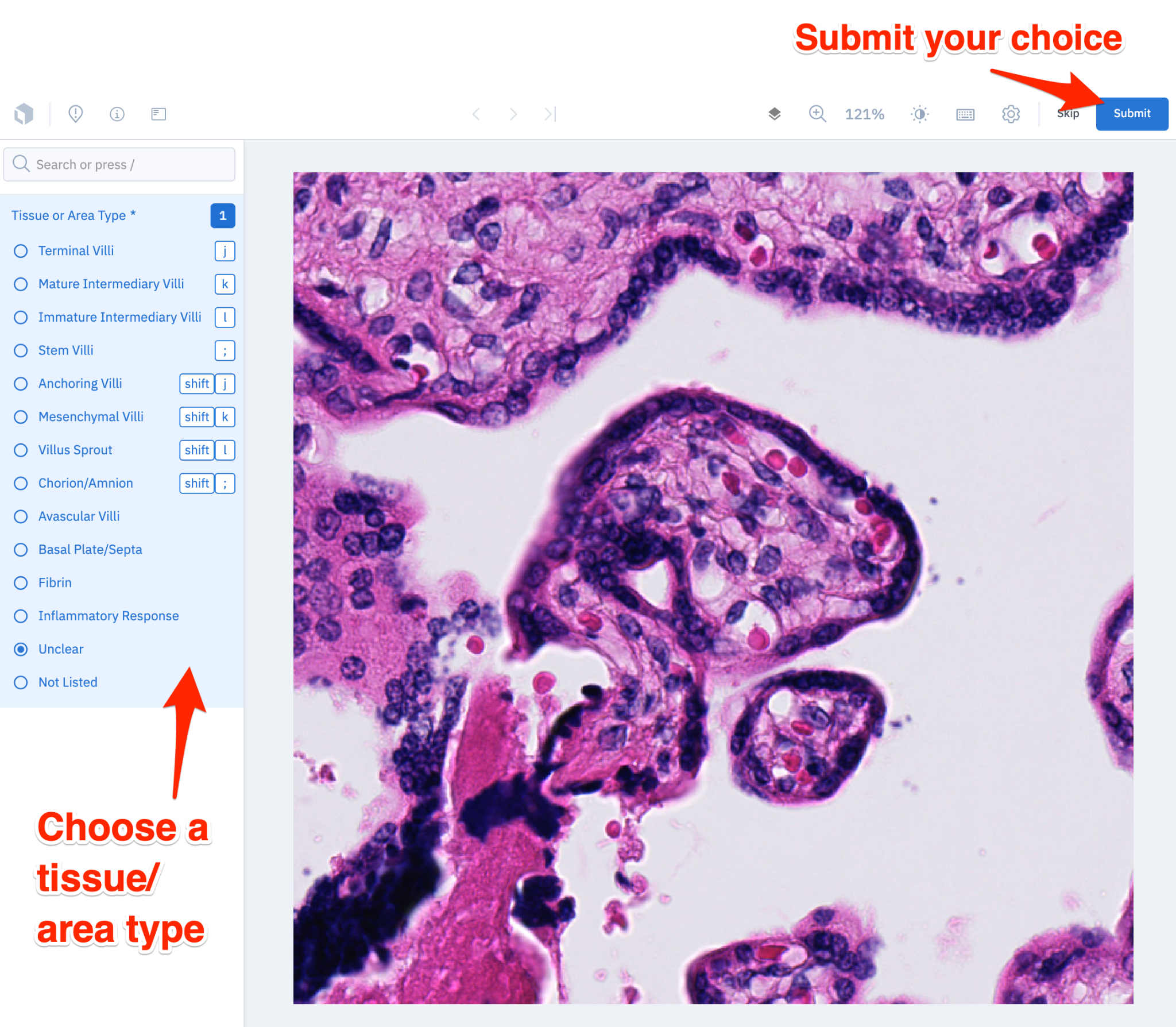

**See Placenta Sample Information**

Press the ‘Info’ icon at the top left to see information about the placenta.

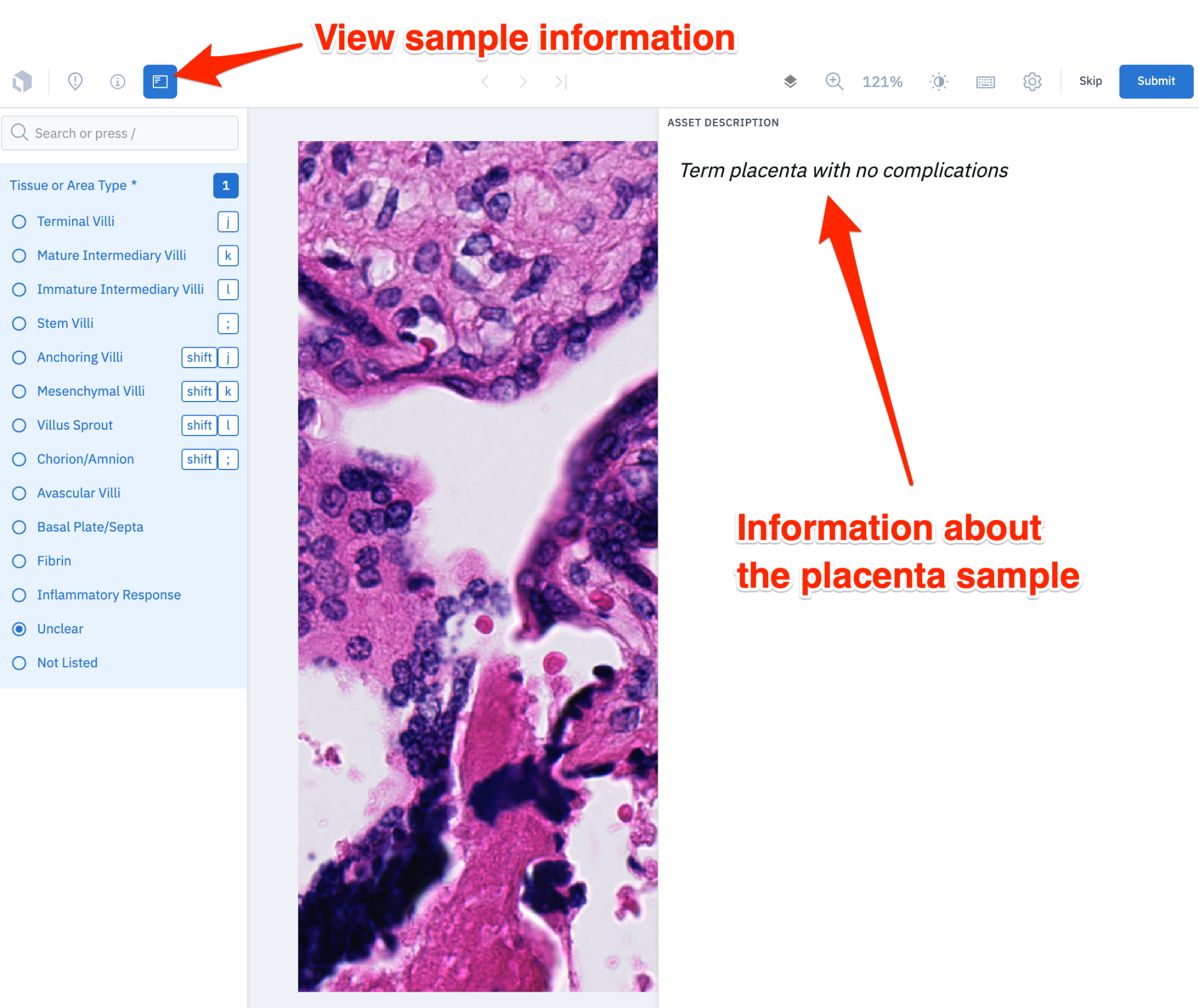

**Add a Comment (Optional)**

Press the ‘Issue’ button at the top left, click on the image, and write a comment in the box on the right.

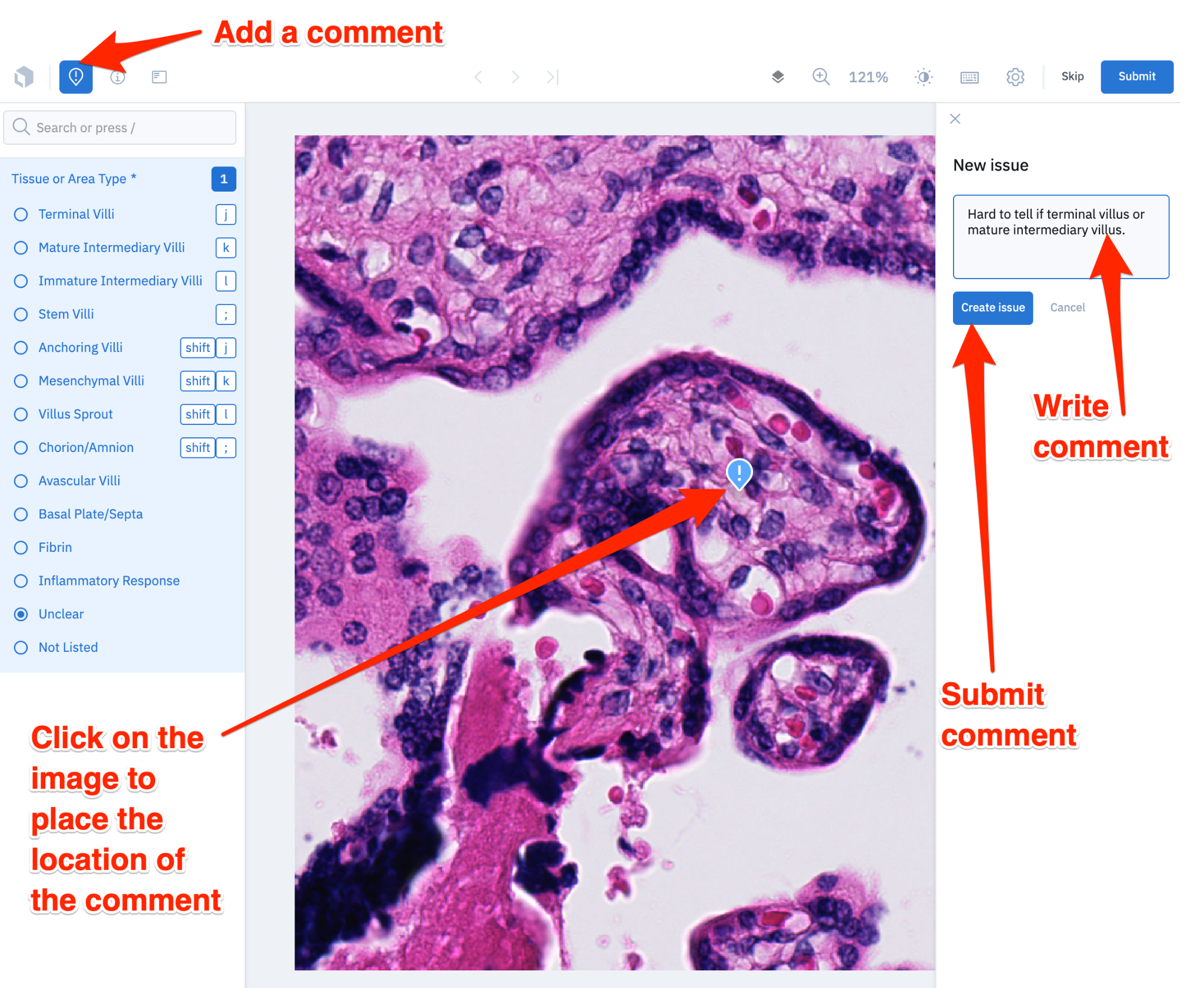

**Tissue Structure Reference Guide**

Literature:

1. <https://link.springer.com/chapter/10.1007/978-3-030-11425-1_36>
   - A comprehensive chapter detailing normal placental development with histology examples and villus tissue diagrams
2. <https://onlinelibrary.wiley.com/doi/10.1111/apm.12858>
   - A summary of placental maturity disorders with histology examples and villus tissue diagrams
3. <https://www.cambridge.org/core/books/placental-and-gestational-pathology/normal-development/11AA85DC1FF68EDA1165A2B46102F1D2>
   - A summary of normal development which focuses on changes across gestational age
4. <https://www.proteinatlas.org/learn/dictionary/normal/placenta>
   - An interactive healthy, term placenta histology slide with some example tissue types
5. <http://eknygos.lsmuni.lt/springer/343/121-173.pdf>
   - A pdf link of Benirschke’s *Architecture of normal villous trees* Chapter from *Pathology of the Human Placenta*
